# Diverse natural variants suppress mutations in hundreds of essential genes

**DOI:** 10.1101/2020.11.24.395855

**Authors:** Leopold Parts, Amandine Batté, Maykel Lopes, Michael W. Yuen, Meredith Laver, Bryan-Joseph San Luis, Jia-Xing Yue, Carles Pons, Elise Eray, Patrick Aloy, Gianni Liti, Jolanda van Leeuwen

## Abstract

The consequence of a mutation can be influenced by the context in which it operates. For example, loss of gene function may be tolerated in one genetic background, but lead to lethality in another. The extent to which mutant phenotypes are malleable, the complexity of the architecture of modifiers, and the identities of causal genes and pathways remain largely unknown. Here, we measure the fitness effects of ~1,500 temperature sensitive alleles of yeast essential genes in the context of variation from ten different natural genetic backgrounds, and map the modifiers for 19 combinations. Altogether, fitness defects for 183 of the 530 tested genes (35%) could be suppressed by standing genetic variation in at least one wild strain. Suppression was generally driven by gain-of-function of a single, strong modifier gene. The validated causes included both variants in protein interaction partners or pathway members suppressing specific genes, as well as general modifiers altering the effect of many temperature sensitive alleles. The emerging frequency of suppression and range of possible suppression mechanisms suggest that a substantial fraction of monogenic diseases could be repressed by modulating other gene products.

## Introduction

The phenotypic outcome of a mutation is determined by the genetic context in which it occurs. The elusive causes of such variation are fascinating in themselves, but are also central to finding ways of predicting and ameliorating genetic diseases. Loss of gene function may lead to death of specific tumour cells only, making the gene a potent drug target (Behan et al. 2019; Gonçalves et al. 2020). Moreover, a coding mutation with no discernible impact in a parent can result in a disorder in their child (Wright et al., 2019). Understanding how such incomplete penetrance arises, and predicting it for a new context, would therefore deepen our understanding of cellular systems, and likely impact diagnoses for developmental disorders or personalised treatments for tumours.

Viability is perhaps the simplest mutation phenotype to analyse. In the course of establishing the yeast gene knockout collection, it became clear that about 1,100 of the ~6,000 yeast genes are indispensable under standard, nutrient-rich growth conditions (Giaever et al. 2002). However, repeating this resource construction in another genetic background offered a tantalising glimpse into the complexity of mutant phenotypes, as over 5% of the essential genes were variable between two closely related strains (Dowell et al. 2010). These and new strain panels (Galardini et al., 2019; Sanchez et al. 2019) have established similar estimates of ~10% of genes demonstrating variable knockout phenotypes between closely related strains and species.

The reason for incomplete penetrance in general, and variable gene essentiality in particular, is the abundance of modifier loci that can suppress mutation effects (Hou et al. 2018). Although their existence has been appreciated for a century (Altenburg and Muller 1920), validated examples remain elusive. A small number of modifiers have been mapped and validated for mouse models (Hamilton and Yu 2012) and human disease (Harper, Nayee, and Topol 2015; Riordan and Nadeau 2017). By far, the most well studied are examples from yeast, powered by the availability of a large number of genetically diverged natural isolates (Peter et al. 2018), genetic tools that allow making large collections of loss-of-function alleles (Hou et al. 2019; Sanchez et al. 2019), and the ability to systematically cross strains in controlled designs (Tong et al. 2001; Hallin et al., 2016; Bloom et al. 2019).

Systematic identification of spontaneous mutations that can suppress fitness defects of “query” mutant alleles in a reference yeast strain has illuminated mechanisms of suppression (van Leeuwen et al. 2016, 2017, 2020). These studies have shown that although deletion mutants are mainly suppressed by genes with a role in the same functional module, partial loss-of-function alleles are frequently suppressed by more general mechanisms affecting query protein expression or stability. However, surveys in model organisms have been largely limited to detecting single gene suppression in a laboratory setting, whereas more complex networks of modifiers may affect the penetrance of any given allele in natural populations. Linkage-based analyses of large panels of individuals have indeed identified second and higher-order modifier effects (Chandler et al. 2014; Taylor and Ehrenreich 2015; Hou et al. 2019; Sanchez et al. 2019), but few modifiers are usually characterised in depth beyond mapping the loci in such designs. The relevance of established broad suppression mechanisms for natural populations thus remains unclear (Matsui, Lee, and Ehrenreich 2017).

Here, we measure phenotypes elicited by crossing nearly 1,500 temperature sensitive mutant alleles of essential genes to ten genetically diverse yeast strains. We use powerful genetic mapping approaches to identify modifier loci of a subset, and validate causal genes for 19 of them. A single strong suppressor allele could independently overcome the mutation phenotype in nearly all mapped cases. The suppressing variants tend to operate within the same biological module as the query gene, with mutations in protein interaction partners or protein complexes often suppressing specific genes, mutations in pathways suppressing other pathway members, and general modifiers altering the effect of many mutations. Together, these results demonstrate the natural genetic flexibility of cells to fulfil crucial tasks, and suggest that loss of human gene function could often be specifically complemented as well.

## Results

### Measuring suppression by standing variation

We set out to test mutation effects in segregant progeny from diverse yeast isolates from various geographic locations and sources (“wild yeasts”) (Liti et al. 2009; Bergström et al. 2014). To do so, we used the Synthetic Genetic Array approach (Tong et al. 2001) to cross a collection of 1,499 temperature sensitive alleles (“TS alleles”) of 673 essential query genes in the laboratory strain S288C (Costanzo et al. 2016) to 10 stable haploid wild yeasts (Figure 1A) (Cubillos, Louis, and Liti 2009), as well as into the S288C control as a reference. We isolated segregant progeny carrying the TS allele to obtain diverse populations of haploid individuals with genomes that, except for the genomic regions around the TS allele and selection markers, are a mosaic of the reference and wild parents (Figure 1B, Methods). We grew the segregants at permissive (26 °C) and restrictive (34 °C) temperatures to measure the fitness effect of TS alleles in different genetic backgrounds. In the control cross with S288C, no segregants are expected to grow at the restrictive temperature due temperature-sensitivity of the allele. However, in cases where the wild yeast strain harbours variants that can suppress the TS phenotype, the haploid segregants that carry them will be able to grow at the restrictive temperature, and will take over the population (Figure 1B).

**Figure 1.**
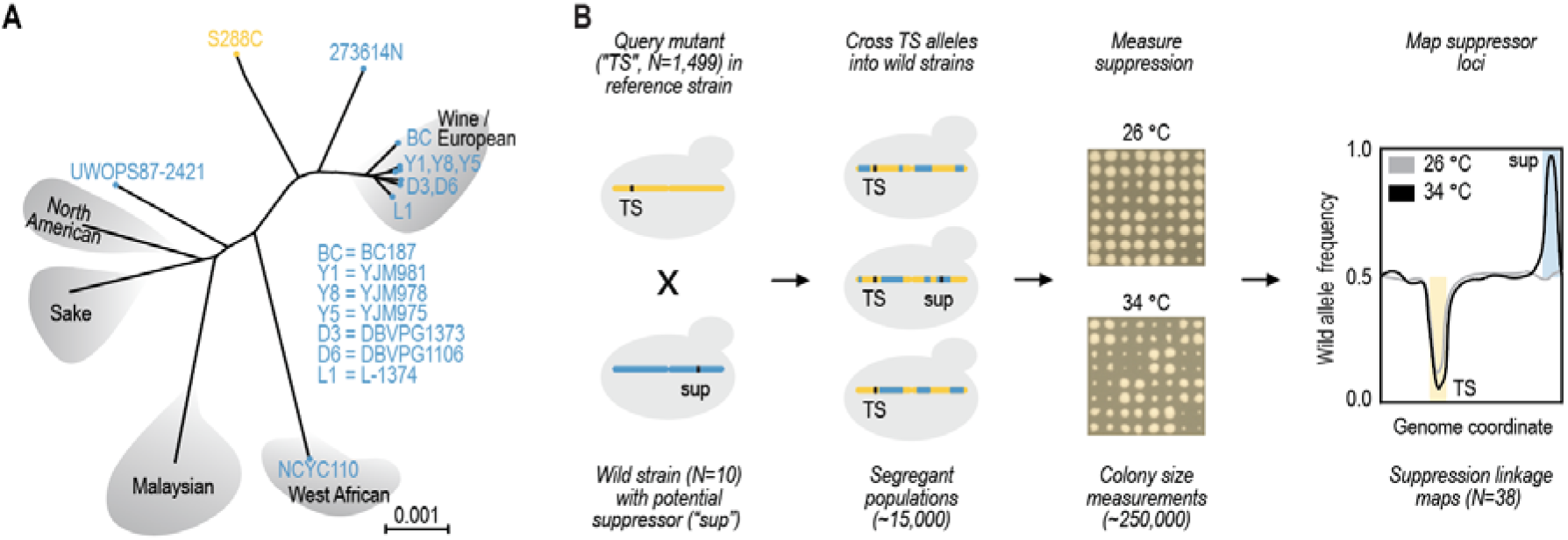
Experimental overview. High-throughput measurement and mapping of suppressor effects from a panel of wild strains. **A**. Strains used in the study. Seven wine / European strains, and three other distant ones (all blue) were crossed to the S288C reference (yellow). **B**. Strategy for identifying suppression by standing variation. A temperature sensitive (TS) allele collection of 1,499 partial loss-of-function mutants in the reference background (yellow) was crossed to the 10 wild yeast strains with potential suppressor alleles (blue), to produce large segregant populations selected to carry the TS allele. The fitness of the resulting 14,990 populations was measured in two biological and four technical replicates. A subset of 38 candidate suppression events were used in bulk segregant analysis for linkage mapping of causal loci that display selection at restrictive temperature (dark) but not permissive temperature (light). Panel A adapted from Liti *et al.* (2009).

We first measured the growth defect of each TS allele in complex pools of wild yeast strain cross progeny (Data S1, Figure S1). After filtering out 379 temperature insensitive strains at 34 °C, we were left with 1,120 TS alleles of 580 query genes (Methods). We estimated suppression as normalised log_2_-scale growth difference between the wild and reference strain crosses at the restrictive temperature, and considered a TS allele phenotype suppressed, if this value was above 0.75, i.e. the wild strain segregants had a 1.68-fold improvement in growth, and if this was unlikely due to chance (false discovery rate = 0.012, Methods, Figure 2A, Data S2).

**Figure 2.**
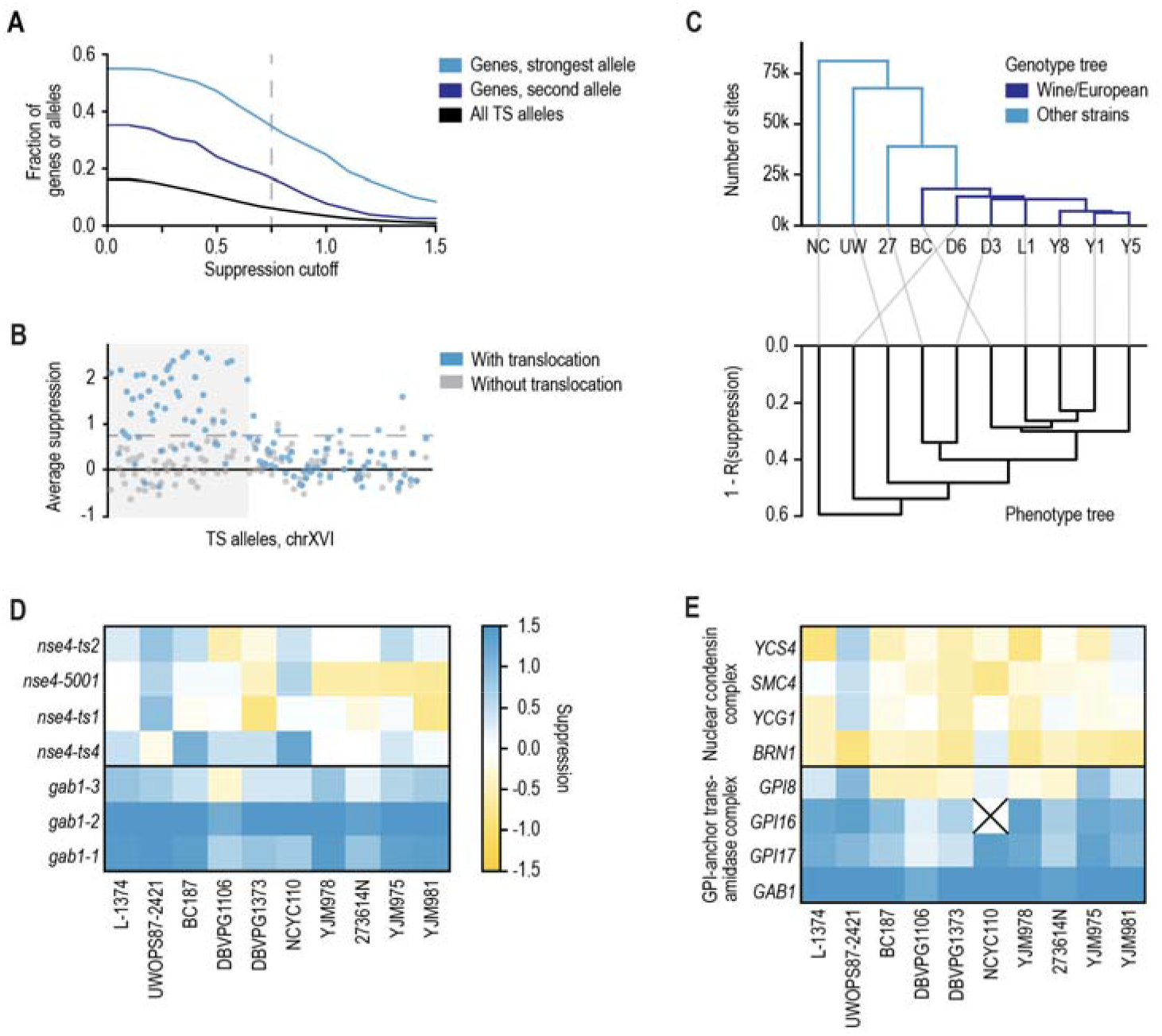
The extent of genetic suppression of essential gene mutants by standing variation. Genetic suppression of TS alleles is not frequent for individual allele-strain combinations, but relatively common for a gene. **A.** Large suppression effects are rare. Fraction of genes or alleles (y-axis) above a suppression cutoff (x-axis) for all pairs of wild strain and allele combinations (black, N=9,708), for the strongest gene suppression signal across alleles and wild strains (cyan, N=529), and for the second-strongest gene suppression signal across alleles and wild strains (blue, N=154). **B.** A positive control region shows suppression signal. Average suppression signal (y-axis) for TS alleles of query genes located on the left arm of chromosome XVI (allele index, sorted by chromosome coordinates, x-axis) across strains with a translocation that generates a duplication of the shaded area (blue markers), and strains without (grey markers). Dashed line: y = 0.75 (suppression cutoff in screen). **C.** Genotype and phenotype trees are concordant. Top: Hierarchical clustering (UPGMA) of the ten wild strains used in this study based on sharing segregating sites. Colours: global genetic cluster membership. Bottom: as top, but based on correlation distance between genetic suppression profiles. Strain abbreviations as in Figure 1A, with in addition: NC = NCYC110, UW = UWOPS87-2421, and 27 = 273614N. **D.** Genetic suppression is consistent across different alleles of the same gene. Suppression score (colour scale) in crosses to different wild strains (x-axis) for TS alleles (y-axis) of *GAB1* and *NSE4* genes. **E**. Genetic suppression is consistent across genes encoding members of the same complex. Strongest suppression score across TS alleles for a gene (colour scale) in crosses to different wild strains (x-axis) for genes (y-axis) that encode members of the GPI-anchor trans-amidase complex (bottom) or the nuclear condensin complex (top). The *GPI16* gene suppression was not estimated in the NCYC110 strain due to chromosome II copy number variation (“X”).

Our screen included several positive control crosses that were expected to show suppression. Three of the wild strains harbour a chrVIII-chrXVI reciprocal translocation (Pérez-Ortín et al. 2002; Figure S2A). Crossing these strains to the reference strain results in 25% of the progeny carrying a duplication of a substantial part of the left arm of chromosome XVI (Figure S2B). This creates an extra copy of the essential query genes in this region that complements the TS allele. Reassuringly, we confirmed that this extra copy suppressed 39 out of 53 TS alleles in the duplicated region on average, while segregant progeny from the other wild strains suppressed a median of two (Figure 2B). Further, the extra copy of chromosome VIII carried by the NCYC110 strain resulted in a similar pattern of suppression (Figure S3A-C).

Also beyond these large genomic determinants, suppression of fitness defects by standing variation in the species was relatively common. Overall, 246 of 1,067 TS alleles (23%, excluding temperature insensitive and copy number suppressed alleles discussed above) and 183 of the 530 tested genes (35%) were suppressed in segregant progeny from at least one genetic background, and on average wild strain progeny could suppress 57 essential gene mutant alleles (5%). For 154 genes, we tested at least two different temperature sensitive alleles, and of these, 25 (16%) had at least two of the alleles consistently showing suppression by some wild yeast variants, with a median of five genes per strain (Figure 2A). Due to variation in temperature sensitivity, different TS alleles of the same gene are not necessarily expected to show suppression at the same temperature. As a negative statistical control, a smaller number of 36 out of 530 successfully tested genes (7%) had at least one allele supporting suppression at the permissive temperature, and 4 of 154 genes with two possible supporting alleles.

To further validate the suppression events identified in our screen, we tested a selection of 102 suppression effects of variable strength by examining the fitness of hundreds of single colony progeny from individual crosses. As the progeny of most crosses showed high variation in colony size, which was also influenced by the number of colonies on the plate, we used stringent thresholds for identifying suppression (Methods). We observed good concordance of strong effects between the phenotypes of the population in the initial screen and the individual progeny in this assay (53% of crosses show suppression for suppression scores above 0.75, 16% for below 0.75, Figure S3E, Data S3).

Thus, we found that suppression by standing genetic variation is relatively common, and that the identified suppression events can often be validated by additional alleles or in complementary assays.

### Patterns of suppression

Next, we asked whether segregant progeny from genetically similar wild strains were more likely to suppress the same TS alleles compared to more diverse strains. Indeed, suppression patterns were more distinct for the two strains genetically furthest from the wine/European cluster (maximum Pearson’s R to any other strain for NCYC110, UWOPS87-2421 less than 0.5, and between 0.59 and 0.75 for the rest), and were consistent with the genetic relatedness otherwise (Figure 2C). The DBVPG1106 wine strain was a phenotypic outlier of Wine/European strains due to overall poor growth at the restrictive temperature (0% of TS allele crosses with log2-scale colony size of more than 10.5 across all crosses; at least 11% for all other strains, Figure S1).

The patterns of suppression of the same gene in the various wild strain crosses were diverse. In some cases, the gene is possibly essential only in the reference background, with multiple alleles showing suppression in all the different tested strains, while for other genes, suppression was limited to a few wild strains (e.g. *GAB1* and *NSE4* genes, respectively, Figure 2D). Suppression was also shared across genes with related function. For example, Gab1 is a member of the GPI-anchor transamidase complex, mutations to three screened members of which were strongly suppressed (Figure 2E), and the *GPI8* gene with less suppression was likely carrying the suppressor variant (see below). Again, we also observed suppression in specific backgrounds, e.g. mutations to genes in the nuclear condensin complex were suppressed almost exclusively in the UWOPS87-2421 background (Figure 2E). In general, we observed consistent suppression patterns between genes encoding members of the same protein complex most frequently (33% of protein complexes with average between-gene suppression correlation significantly higher than permuted controls at FDR=0.20, Methods), followed by cellular locations (30%), broad functional categories (28%), KEGG pathways (27%) and Gene Ontology categories (25%). This concordance is consistent with the nature of connectedness within genetic networks in general, where many interactions are shared within complexes, compared to broader functional connections.

### Mapping of genomic regions involved in suppression

Given frequent, strong, and technically and biologically consistent suppression of TS alleles by variants from wild genetic backgrounds, we next sought to identify the causal loci and genes. First, to estimate the average number of modifiers involved in the suppression phenotype, we dissected meiotic progeny of 16 crosses, and examined the growth of spores carrying the TS allele at 26 and 34 °C. In all cases, 15-65% of the spores grew well at the restrictive temperature, with little additional phenotypic variation in growth beyond survival, suggesting that most of the detected suppression phenotypes are the result of at most 1-3 strong modifier variants in the wild strain background (Figure S4, Discussion).

To map the suppressor loci, we performed bulk segregant analysis on 38 segregating TS allele populations at both 26 °C (allele functional) and 34 °C (allele loss-of-function, Figure 1B) (Liti and Louis 2012). We sequenced the populations, and compared variant allele frequencies between the two temperatures (Data S4 and S5). We first considered positive controls expected to involve suppression by an additional, wild-type copy of the query gene described above, either generated by the chrVIII-chrXVI translocation (6 samples) or located on an aneuploid chromosome (6 samples). In all 12 cases, we could indeed observe selection for either the translocation or the aneuploidy, and further confirmed that suppression occurred by the presence of a second, wild-type allele of the query gene (Figures S2C-D and S3D).

Second, we sequenced meiotic progeny of six crosses that showed weak “suppression” in our screen (suppression score below 0.6), unrelated to any known translocations or aneuploidies. Four cases showed selection for newly acquired aneuploidies of either the chromosome carrying the query gene or other loci selected in the crossing protocol. These cases often represent ways of cells to escape the strong selection applied in our protocol, rather than true cases of suppression. The remaining two cases harboured regions of selection for the wild strain sequence specific to high temperature (Data S4 and S5), suggesting that some of the weaker scores in our screen also represent true cases of suppression, and corroborating the observations from the confirmation of individual suppression effects.

Third, we analysed 20 crosses that showed strong suppression in our screen (suppression score above 0.75). The large majority (16) showed regions of specific selection for the wild sequence at high temperature (Figure 3A, Data S4 and S5), whereas three populations diploidised or showed selection for an aneuploidy. The one remaining cross did not show any suppressor loci or aneuploidies. Thus, we could map suppressor loci for 16/20 (80%) of the crosses that showed strong suppression in our screen, and in 2/6 (33%) of the crosses that showed weak suppression.

**Figure 3.**
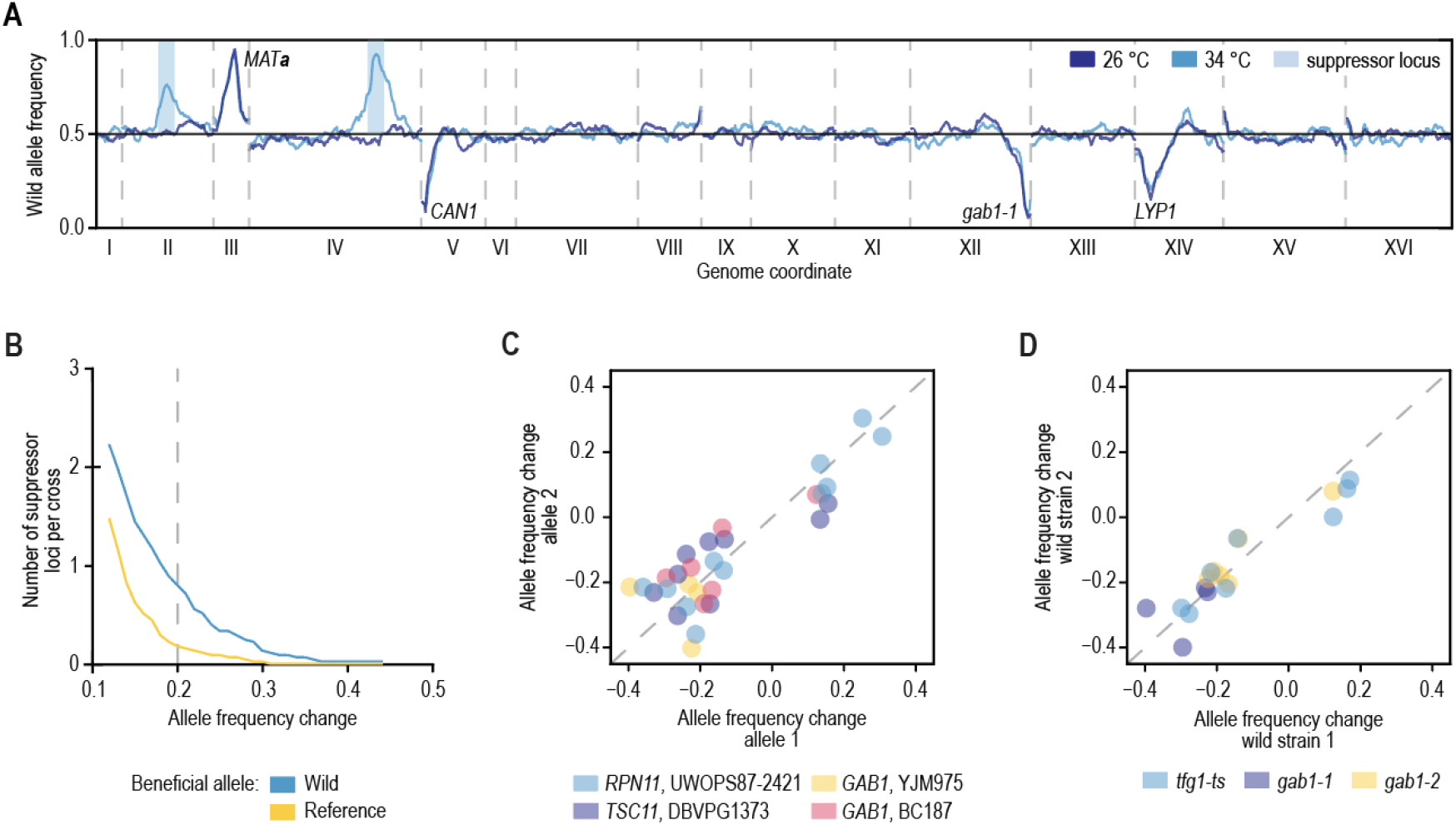
Mapping suppressor loci. Suppressor loci can be mapped by sequencing segregant pools. **A.** Example of the mapping results. Wild allele frequency in progeny of a cross of the *gab1-1* temperature sensitive allele to the YJM975 strain (y-axis) along the yeast genome (x-axis) at the permissive 26 °C (blue) and restrictive 34 °C (allele loss-of-function, cyan). The allele frequency change between the two temperatures is used in mapping. Labels: selected loci in the cross. Blue regions: called suppressor loci. **B.** Suppressors are plentiful. The average number of suppressor loci per cross (y-axis) at given allele frequency change cutoff (x-axis) with wild allele beneficial (blue) or reference allele beneficial (yellow). **C.** Suppressors are reproducible across TS alleles. Allele frequency change in crosses using different TS alleles of the same gene (x- and y-axis) crossed with the same wild strain. Colours: gene and wild strain combinations. **D**. Suppressors are reproducible across wild strains. Allele frequency change at 34 °C in crosses using the same TS allele and different wild strain (x- and y-axis). Colours: TS alleles.

The landscape of suppressors is diverse. We identified 31 suppressor loci in the 19 crosses without aneuploidies (1.6 on average, Figures 3B and 4A, Data S5), and an additional 48 weaker reproducible signals (2.5 on average, Figures 3B and 4A, Data S5). This number of modifier loci is in agreement with our estimates based on segregation patterns observed after tetrad dissection (Figure S4). Most of the suppressor loci (27/31) were selected for the wild strain sequence, consistent with the additional variation in species providing the substrate for circumventing essential gene function. Reassuringly, suppressor loci were reproducible across biological replicates, different TS alleles of the same gene when crossed to the same wild strain (e.g. *RPN11*), and same TS alleles when crossed to different wild strains (*TFG1* and *GAB1;* Figures 3C-D and 4A). A suppressor locus on chromosome XIV was shared across five different essential genes (*GPI13*, *MED7*, *RPN11*, *SEC24*, and *TFG1*), indicating the presence of a pleiotropic modifier in this locus, co-localising with the previously characterised *MKT1* gene (Steinmetz et al. 2002).

### Suppressor gene identification and validation

We next sought to identify the causal suppressor genes. As each of the mapped suppressor regions harbors tens of plausible candidate genes, we computationally prioritised them based on their functional connection to the query gene (Data S6, Methods, (van Leeuwen et al. 2020)). We also included known general modifiers, such as *MKT1* and *HAP1*, that each affect the expression of thousands of genes (Fay 2013; Parts et al. 2014; Albert et al. 2014, 2018). To test the phenotypic consequence of the candidates, we replaced their open reading frame and ~100-400 bp surrounding region with the wild version in the reference strain background, and tested for suppression of the corresponding TS allele. In total, we tested 50 suppressor gene candidates from 31 mapped loci of various strength (Data S7, Figure S5), and identified causal genes for 17 loci (55%, Data S7, Figures 4A and S5), validating both our mapping strategy, as well as the computational prioritisation. The 14 suppressor loci without a confirmed suppressor gene resulted from failed experiments, inconclusive results, or cases in which no suppression was observed for the wild allele (Data S7). Causal suppressor genes were more likely to be identified for strong suppressor loci compared to weaker signals, suggesting that in unconfirmed cases the suppression phenotype may have been too weak to detect in our validation assay (Figure 4B).

**Figure 4.**
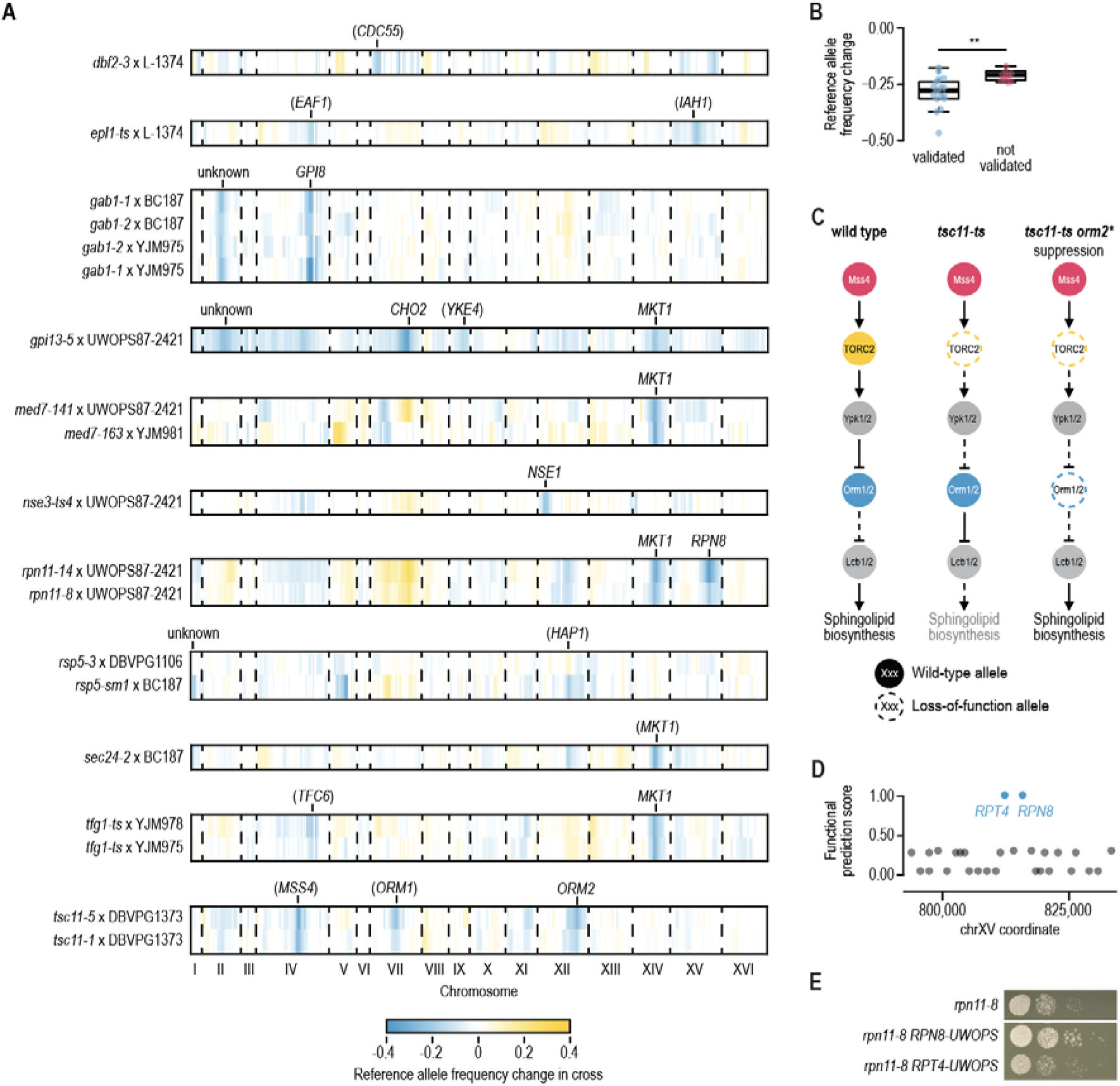
Identifying and validating suppressor candidates. **A.** Mapping results for segregant pools involving the indicated TS alleles and wild strains. The change in S288C allele frequency between the meiotic progeny isolated at 26 °C and 34 °C is plotted by genomic coordinate. Causal suppressor genes are indicated for regions that show selection for the wild strain sequence. Genes in brackets have not been validated experimentally. **B.** Comparison of the change in reference allele frequency, either for suppressor loci for which a causal suppressor gene was validated (N=17), or for loci for which we were unable to validate a suppressor gene (N=10). Loci for which all experiments failed due to technical reasons were excluded from the analysis. Statistical significance was determined using a two-tailed Mann-Whitney’s U test (** = p < 0.005). **C.** A cartoon of the TORC2 signalling pathway, highlighting suppression of a TORC2 mutant (*tsc11*) by mutations in *ORM2*. **D.** Suppressor prediction within the chromosome XV QTL of *rpn11* TS mutants. Functional information prioritization score (y-axis) for genes in the suppressor region (x-axis) identified *RPT4* and *RPN8* as two highest-scoring suppressor candidates. **E**. Experimental validation of *RPN8* as the causal suppressor of the *rpn11-8* TS mutant. Cultures of the indicated strains were diluted to an optical density at 600 nm of 0.1 and a series of ten-fold dilutions was spotted on agar plates and incubated for 2-3 days at 34 °C. UWOPS = UWOPS87-2421. See also Figure S5.

Many of the validated suppressor genes were consistent with their known roles in the biology of the query gene. For example, two different alleles of *TSC11*, which encodes a subunit of TOR complex 2 (TORC2) that activates a phosphorylation cascade controlling sphingolipid biosynthesis, were suppressed by multiple variants present in the DBVPG1373 strain (Figure 4A). The two strongest suppressor loci were located around *MSS4* and *ORM2*, which encode an upstream activator and a member of the TORC2 signalling pathway, respectively (Lucena et al., 2018; Han et al., 2010; Figure 4A and C). Indeed, we confirmed the suppression of *tsc11-5* temperature sensitivity by the *ORM2-DBVPG1373* allele (Figure S5). Orm2 inhibits the first committed step in sphingolipid synthesis, and loss of Orm2 function may lead to the reactivation of sphingolipid biosynthesis in the absence of TORC2 (Figure 4C). Intriguingly, a third weak suppressor locus was identified for the *tsc11-5* allele around *ORM1*, a paralog of *ORM2*. However, *ORM1* has no variants within the ORF in the DBVPG1373 strain, and we were not able to confirm a suppression phenotype for the *ORM1* promoter variants in the presence of reference alleles of *MSS4* and *ORM2*.

Genetic mapping alone is not sufficient to identify causal genes. Our computational prioritisation identified *RPT4* and *RPN8* as potential suppressor genes within the chromosome XV suppressor locus of *RPN11* with equally high scores (Figure 4A and D). As the query gene *RPN11* encodes a metalloprotease subunit of the 19S regulatory particle of the proteasome, and the candidate suppressors *RPT4* and *RPN8* also both encode subunits of the same particle, genetic information and computational prior were not sufficient to pinpoint one as the causal gene. In experimental validations, the *RPN8* allele from UWOPS87-2421 suppressed the *rpn11-8* phenotype, whereas the *RPT4* allele did not (Figures 4E and S5). Rpn8 and Rpn11 form an obligate heterodimer (Bard et al. 2018), and the *RPN8* allele from UWOPS87-2421 may thus restore the interaction between the two proteins, which could have been weakened by the *RPN11* mutations. This ability to resolve a causal gene from multiple linked candidates underscores the importance of thorough experimental validation to understand the mechanism of suppression.

### Genetic simplicity of strong suppression

Previous approaches for identifying suppressors have relied on spontaneous mutation, and thus sample genetic backgrounds that are very similar to that of the reference. As a result, more complex allele arrangements that may be required for suppression, e.g. combinations of two or more mutations, are not easily obtained. Despite observing multiple loci that are involved in the suppression phenotype in each of the sequenced populations (Figure 4A), we found no evidence for the interdependence of one suppressor locus genotype on the presence of another, and all strong suppressors acted in isolation (Figures 4B and S5; Discussion). For example, both the *RPN8* and the *MKT1* allele from UWOPS87-2421 could individually suppress the *rpn11-14* TS allele to near-wild type fitness (Figure S5). We did not observe examples consistent with strong suppression by many small effect variants. Conversely, multiple mutations within a locus could be required for robust suppression. The *ORM2-DBVPG1373* allele that independently suppressed *tsc11-5* carries two missense mutations, P26T and G134S, that affect conserved residues, are predicted to be highly deleterious, and are both required for a robust suppression phenotype (Figure S5).

Next, we compared the suppressor genes identified by our mapping results to those previously found in the reference strain background (Oughtred et al. 2019). In cases where we confirmed a candidate suppressor, we also often found prior evidence of suppression in that gene (4/9 unique suppressor-query pairs; Data S7). In all four cases, the suppressor and query gene pairs encoded members of the same protein complex or pathway, and in three cases the suppressor and query proteins interact physically (Data S7). The five suppressor-query gene pairs that had not been previously described included four cases of suppression by the general modifier *MKT1*. We have previously observed a similar prevalence of general suppressor genes that affect the expression of the query mutant among spontaneous suppressor mutations of TS alleles isolated in the reference strain (~50% of all suppressor genes; (van Leeuwen et al. 2016, 2017)). Out of the 9 suppressor-query pairs, 7 (78%) appear to involve a gain-of-function suppressor allele (Methods; Data S7). This relatively large proportion of gain-of-function alleles is consistent with the idea that loss-of-function alleles may be under stronger negative selection in natural populations.

Overall, mechanisms of suppression identified in a laboratory setting mimic those driven by natural variation, and can involve identical suppressor genes when considering suppressors that function within the same functional module as the query gene. In addition, despite the presence of multiple selected suppressor regions in nearly every cross, strong suppressor mutations always acted independently of the genetic background. Combined, these observations are consistent with a model where single genes evolve along a lineage, perhaps adapting to the rest of the environmental and genetic context via multiple gain-of-function mutations, which then in turn gives the derived allele ability to independently suppress fitness effects of other alleles.

### Mutations in NSE1 can suppress SMC5/6 complex dysfunction

One of our mapped suppressor interactions involved the suppression of a *nse3-ts4* TS allele by the *NSE1* allele from UWOPS87-2421 (“*NSE1-UW*”; Figures 4A and 5A). Nse1, Nse3, and Nse4 form a subcomplex within the highly conserved SMC5/6 complex, which is essential for the removal of recombination intermediates during DNA replication and repair (Figure 5B) (De Piccoli et al. 2006; Menolfi et al. 2015). Nse3 and Nse4 bridge the globular head domains of Smc5 and Smc6 (Figure 5B), whereas Nse1 is a RING finger protein with ubiquitin ligase activity that strengthens the interactions between Nse3 and Nse4 (Hudson et al. 2011; Pebernard et al. 2008). The *NSE1-UW* allele also suppressed the growth defect of a *nse4-ts4* TS allele, but not that of any of the other tested SMC5/6 subunits (Figure 5A). A *nse1* loss-of-function allele exacerbates the fitness defect of a *nse3* TS mutant (Costanzo et al. 2016), and overexpression of *NSE1* suppresses a *nse3* TS allele (Magtanong et al. 2011) (Figure 5C), suggesting that the *NSE1-UW* allele has a gain-of-function effect that improves the stability or activity of the SMC5/6 complex. Indeed, the *NSE1-UW* allele suppressed the sensitivity of *nse3* and *nse4* mutants to DNA damaging aging agents hydroxyurea (HU) and methyl methanesulfonate (MMS) (Figures 5D and S6A), but could not suppress the lethality associated with deleting either *NSE3* or *NSE4* (Figure S6B). Thus, SMC5/6 complex activity could be restored in *nse3* and *nse4* partial loss-of-function mutants by the presence of *NSE1-UW*.

**Figure 5.**
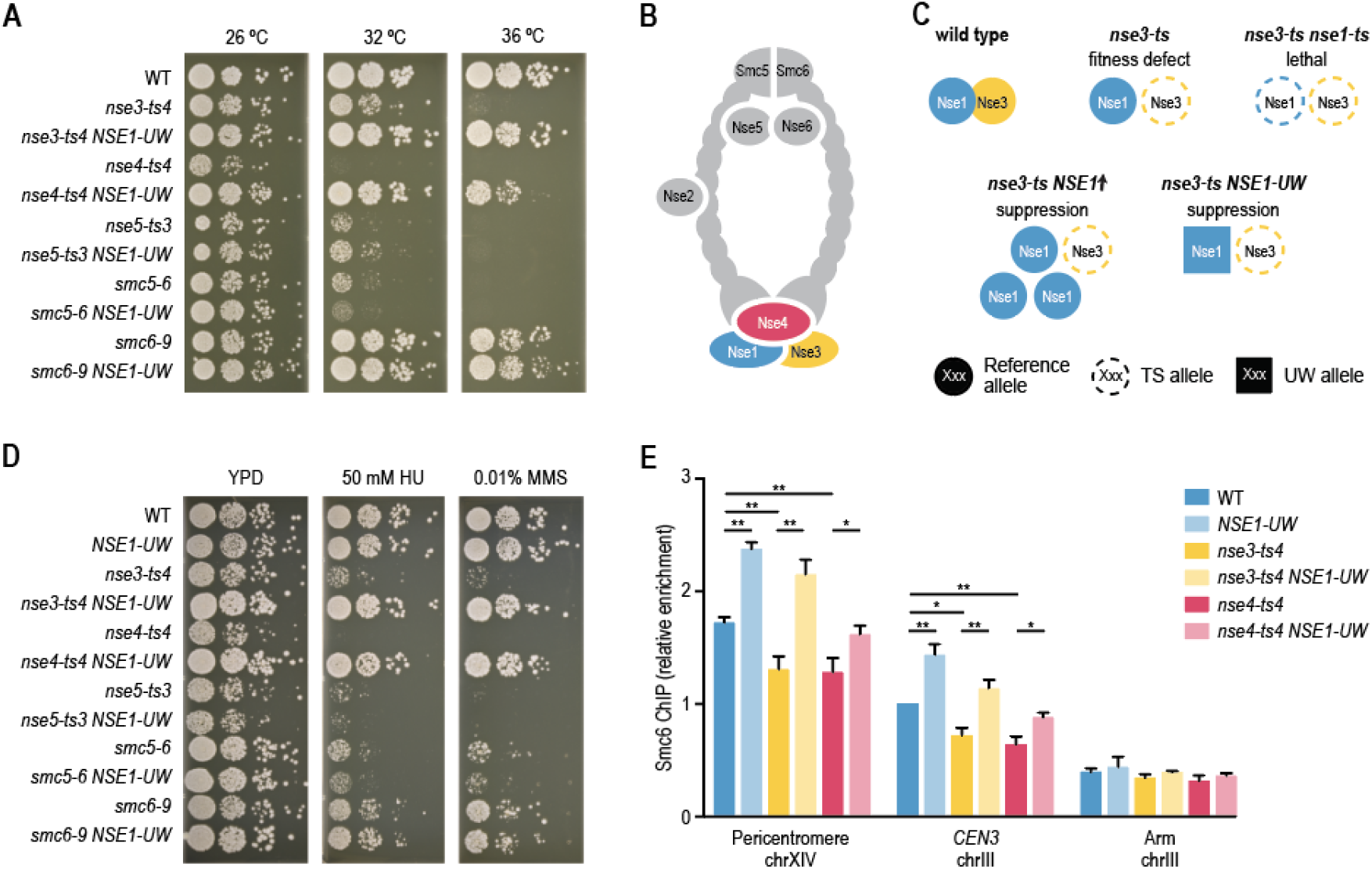
The *NSE1* allele of UWOPS87-2421 can suppress *NSE3* and *NSE4* TS mutants. **A,D.** Suppression of *nse3-ts4* and *nse4-ts4* temperature sensitivity (A) and DNA damage sensitivity (D) by the *NSE1* allele of UWOPS87-2421. Cultures of the indicated strains were diluted to an optical density at 600 nm of 0.1 and a series of ten-fold dilutions was spotted on agar plates and incubated for 2-3 days. UW = UWOPS87-2421. The plates shown in (D) were incubated at 30 °C. **B**. A cartoon of the SMC5/6 complex. **C**. An illustration of the various types of genetic interactions that have been observed between different alleles of *NSE1* and *NSE3*. **D**. See (A). **E**. Recruitment of Smc6-FLAG by ChIP-qPCR at two known SMC5/6 binding sites (pericentromere of chromosome XIV and *CEN3*) and one negative control locus (arm of chromosome III) in G2/M arrested strains. Relative enrichment corresponds to the ratio of the signal after immunoprecipitation (FLAG) over beads alone, normalized to the WT ratio at *CEN3*. Error bars: standard error of the mean of three independent experiments. Statistical significance was determined using Student’s t-test (* = p < 0.05, ** = p < 0.005).

To more directly test the impact of *NSE1-UW* allele on the SMC5/6 complex function, we measured its accumulation at two established chromosomal SMC5/6 binding sites using an Smc6-FLAG based ChIP-qPCR assay in the reference strain (Lindroos et al. 2006; Jeppsson et al. 2014). Although the amount of Smc6-FLAG protein was comparable in all strains (Figure S6C), the accumulation of Smc6-FLAG at the two genomic loci was substantially reduced in *nse3-ts4* and *nse4-ts4* mutants compared to wild type (Figure 5E). Replacing the reference *NSE1* allele with the *NSE1* allele of UWOPS87-2421 increased recruitment of the SMC5/6 complex to the DNA, both in the presence of wild-type or TS alleles of *NSE3* or *NSE4.* This suggests that the *NSE1-UW* allele increases association of the SMC5/6 complex to the DNA, thereby counteracting the negative effects of the *nse3* and *nse4* TS alleles on SMC5/6 complex activity.

## Discussion

We used systematic large-scale genetics to cross partial loss-of-function alleles to ten different genetic backgrounds, measured the extent to which standing variation in the species can suppress the loss-of-function phenotype, used a powerful pooled mapping approach to localise modifying alleles, and identified the causal genes for many of them. We found that suppression was consistent across replicates, alleles, and within complexes, with modifier genes often acting either directly by interacting with the mutated protein or the complex in which it operates, complementing the output of the pathway in which it is a member, or unspecifically via general compensation mechanisms.

### Genetic architecture of suppression

The genetic architecture of suppression we mapped was skewed towards alleles of strong effect. Nearly all linkage maps had a strong modifier, and there was a long tail of weaker effects, many of which validated, as has been observed for virtually all mapped traits in general. Some previous studies identified a relatively large fraction of beneficial reference alleles in yeast linkage maps (e.g. ⅓ of overall in Parts et al., 2011). Although we find a comparable fraction of reference alleles when including weak modifier loci, we found the reference allele was preferred in only every eighth strong suppressor locus. This is consistent with a few large effect alleles explaining the majority of phenotypic variability, that then by definition requires their effect to align with the phenotype differences between the strains.

We attempted to identify candidate genes for all strong suppressors, and succeeded for most. Multiple TS alleles and different wild strains often consistently supported the suppression region, and the plausible suppression mechanisms were affected via gain-of-function mutations affecting complex integrity, pathway activity, or unspecific modifiers. For the three strong suppressor loci for which we failed to confirm a suppressor candidate (Figure 4A), we may have failed to include the causal suppressor gene in our experiments, or the suppression may have been dependent on the presence of other suppressor alleles. However, cases of suppression that require more than a single allele were not frequent in our analysis, as a single strong suppressor allele could independently overcome the mutation phenotype in nearly all mapped cases. These results are consistent with a long list of studies that identify a single gene or genomic locus with a strong effect on a phenotypic trait in diverse organisms (Johnston et al. 2011; Barson et al. 2015; Jones et al. 2018; Thompson et al. 2020), and suggest additional scrutiny of single large effect alleles modifying human phenotypes as a promising research direction as well.

Many of the suppression events were driven by aneuploidies. These generally involved pre-existing aneuploidies and translocations, most frequently in the wild parent. This is not surprising, as the wild strains generally tolerate aneuploidy well (Hose et al. 2015; Peter et al. 2018), and the strong imposed selection forces the cells to use all available diversity to survive. The range of possible ways to escape the various selection steps was such that it is arguable that most of the logically consistent and physically possible scenarios took place. While such chromosome-scale plasticity may not be common in higher eukaryotes where imbalances in gene dosage are often deleterious, it underscores the evolutionary potential of large-scale rearrangements compared to point mutations. *De novo* aneuploidies and diploidisation events also explain a large fraction (67%) of the weak “suppression” signals that we sequenced, with a suppression score <0.75 in our screen, where a few cells had escaped one of the selection steps, and could partially take over the population.

### Consistency of suppression across experiments

A subset of the genes we mapped suppressors for had previously been analysed to identify spontaneously evolved modifiers in the reference background. The suppressor genes that had a functional connection to the query gene were often identical in both studies, consistent with the shared selection targets of *de novo* and pre-existing variation observed under drug treatment (Li et al. 2019). This could indicate that the suppressor allele complements an independent deficiency in the reference strain (consistent suppression of *GAB1* in all wild strains by the same *GPI8* allele), or that the suppressor has co-evolved with the complex or pathway within (SMC5/6 complex and TORC2 pathway). Further, the vast majority of validated suppressor alleles likely conferred a gain-of-function phenotype compared to the reference allele (Data S7), whereas many of the spontaneous suppressor mutations isolated in the reference background had a loss-of-function effect (van Leeuwen et al. 2016, 2020). Loss-of-function alleles are more likely to arise spontaneously as the underlying mutation events are more common, but may have a higher chance to be subjected to negative selection in natural population compared to gain-of-function variants.

We also frequently observed suppression via general, pleiotropic modifiers. Although general modifiers that can suppress the growth defect of many different mutant genes have been identified by spontaneous mutation in the reference background as well, they tend to affect mRNA and protein degradation pathways (van Leeuwen et al. 2016, 2017). The natural variation general modifiers *HAP1*, encoding a transcription factor regulating the response to haem and oxygen, and *MKT1*, encoding a nuclease-like protein of which the precise cellular function remains unclear, were never found as spontaneous suppressors among the >2000 described suppressor interactions in S288C (van Leeuwen et al. 2016). In the case of *HAP1* this is expected as the gene is inactivated by a transposon insertion in S288C. The *MKT1* gene on the other hand is intact in S288C, but the reference allele may perform poorly compared to *MKT1* alleles available in the wild, that have been described to suppress many different phenotypes, including temperature sensitivity (Steinmetz et al. 2002; Fay 2013; Parts et al. 2014; Albert et al. 2014, 2018).

Our study used temperature sensitive mutant strains that show a progressive decline in gene function with an increase in temperature. This enables identifying suppressors that can completely bypass gene function, but also those that rescue partially functional alleles. For example, the *nse3-ts4* allele could be completely rescued by mutations in *NSE1*, both encoding members of the Nse1-Nse3-Nse4 complex module (Figure 5). However, this subcomplex would not assemble in the absence of the *NSE3* gene, and the *NSE1* mutant allele does not rescue a nse3Δ deletion mutant (Figure S6). Comparison of our suppressors with a systematic survey of bypass suppression of essential gene deletion mutants (van Leeuwen et al. 2020), showed little overlap in the identified suppressable essential genes, suggesting that the vast majority of standing variation suppressors will depend on the presence of the TS allele.

### Future perspectives

Our screen for natural variants that can suppress TS alleles was not saturated. First, although the TS mutant strain collection we used in our screen contained TS mutants for ~60% of all essential yeast genes, due to variation in temperature sensitivity, not all tested genes will have had a suppressable phenotype at our chosen restrictive temperature of 34°C. Second, the set of possible suppressor mutations we considered was restricted to the standing variation in the ten wild strains we used. Indeed, we could not detect all known suppression alleles that have been identified via spontaneous mutation in the reference background. Despite these limitations, we found that 35% of the tested essential genes could be suppressed by at least one wild strain. As this relatively high number is likely an underestimate of the true suppression potential of standing variation, we expect suppression to be common in natural populations.

We have provided a first glimpse into the extent, complexity, and mechanisms of mutation effect suppression by standing variation. Given the high frequency at which we observed suppression via complementing natural variants, we expect it to have an important contribution to other phenotypes, species, and contexts, including human disease. The large overlap between natural suppressor variants and those identified in a laboratory setting suggests that suppressor screening in human cell lines will help understand variable penetrance of human disease mutations as well. In parallel, systematic studies in yeast and other species will continue to refine our view of the mechanisms adopted by modifier mutations to determine the severity of genetic traits.

## Supplementary Data

1. Raw colony size data
2. Suppression scores
3. Random sporulation analysis
4. Allele frequency VCF files
5. Identified QTLs
6. Suppressor gene prediction
7. Tested suppressor gene candidates
8. Yeast strains and plasmids

## Acknowledgements

We thank M. Costanzo, J. Hou, S. Soyk, G. Tan, M. Taschner, B. Ünlü, and A. Vjestica for critical reading of the manuscript, discussions, reagents, and technical assistance. We also thank B. Andrews and C. Boone whose lab the work was incepted in, and whose vision shaped the direction of this project. L.P. was supported by Wellcome (206194), IT Centre of Excellence EXCITE (TK148), and a Marie Curie International Outgoing Fellowship (328541). J.v.L. was supported by the Swiss National Science Foundation (PCEGP3_181242). G.L was supported by the Foundation for Medical Research (EQU202003010413). C.P. holds a Ramon y Cajal fellowship (RYC-2017-22959). The authors declare that they have no conflicts of interest.

## Methods

### Yeast strains, plasmids, and growth

Yeast strains were grown using standard rich (YPD) or minimal (SD) media. Methyl methanesulfonate (MMS) and hydroxyurea (HU) were obtained from Sigma-Aldrich.

For SGA analysis (see below), we used a collection of temperature sensitive mutants of essential genes (*MATα xxx-ts::natMX4 can1Δ::STE2pr-SpHIS5 lyp1Δ his3Δ1 leu2Δ0 ura3Δ0 met15Δ0*; (Costanzo et al. 2016)). Four of these strains appeared to have a different TS mutant allele than originally annotated. Because we could not determine where a potential mistake or mix-up had occurred, we assigned new strain IDs to these strains. TSQ2353 (*tre2-5008*) was renamed as TSQ2884x (*tfg1*), TSQ1864 (*brr2-5019*) as TSQ2885x (*fas2*), TSQ1877 (*iki3-5008*) as TSQ2886x (*epl1*), and TSQ1879 (*iki3-5010*) as TSQ2887x (*epl1*).

For the allele swaps (see section “suppressor candidate validation”) we used strains from either the BY4741 deletion mutant collection (*MAT****a*** *xxxΔ::kanMX4 his3Δ1 leu2Δ0 ura3Δ0 met15Δ0*; Euroscarf), or the TS-allele-on-plasmid collection (*MAT****a*** *xxxΔ::natR_kanR(Cterm) his3Δ1 leu2Δ0 ura3Δ0 [xxx-ts_kanR(Nterm), AgMFA2pr-hphR, URA3]*; (van Leeuwen et al. 2020)).

All other yeast strains used in this study are listed in Data S8.

### Making the wild yeast strains SGA compatible

Twenty-six wild yeast strains had previously been deleted for *HO* and *URA3*, and haploid *MAT****a*** spores had been isolated (*MAT****a*** *hoΔ::hphMX6 ura3Δ::kanMX4*; (Cubillos, Louis, and Liti 2009)). To make these strains compatible with SGA analysis and facilitate further genetic manipulations, we (partially) deleted the *LEU2* and *HIS3* genes.

First, to delete *LEU2*, we used plasmid p7410 (Data S8), that contains in the following order: a SwaI restriction site, base pair −403 to 8 of *LEU2*, base pair +62 to +258 downstream of the *LEU2* stop codon, the *TDH3* promoter from Ashbya gossypii (Ag) driving the *nrsR* (“*natR*”) gene followed by the *AgTDH3* terminator, the *GAL1* promoter driving *KAR1* followed by the *AgCYC1* terminator, the *kanMX4* cassette, and base pair +62 to +783 downstream of the *LEU2* stop codon. We digested the plasmid using SwaI, and transformed the wild yeast strains with the linearised plasmid. Transformants were isolated on YPD + NAT, and subsequently replica plated onto YPGal media, to induce overexpression of *KAR1*, which is lethal and thus selects for recombination between the two *LEU2-3’* sequences. Recombination was confirmed by testing for growth on YPD + G418 media.

Second, to partially delete *HIS3*, we used plasmid p7411 (Data S8), that contains in the following order: a SwaI restriction site, base pair 137 to 310 of the *HIS3* gene, base pair 495 to +112 of the *HIS3* gene, the *AgTEF1* promoter driving *LEU2* followed by its endogenous terminator, the *URA3* gene under control of its own promoter and terminator, and base pair 495 to +707 bp of *HIS3*. We digested the plasmid using SwaI, and transformed the wild yeast strains with the linearised plasmid. Transformants were isolated on SD -Ura -Leu, and subsequently replica plated onto media containing 5-fluoroorotic acid (SD + 5-FOA), which is toxic to cells expressing *URA3* and will thus select for recombination between the two *HIS3-3’* sequences. Recombination was confirmed by testing for growth on SD -Leu media.

Proper deletion of *LEU2* and a part of *HIS3* was confirmed by PCR. Strain identity was validated by sequencing the barcodes inserted at the *ura3Δ* locus (Cubillos, Louis, and Liti 2009). In total, we obtained 10 wild yeast strains with the genotype *MAT****a*** *hoΔ::hphMX6 ura3Δ::kanMX4 his3Δ1 leu2Δ0* (Data S8).

### Synthetic genetic array (SGA) analysis

The 10 SGA-compatible wild strains (Data S8, *MAT****a*** *hoΔ::hphMX6 ura3Δ::kanMX4 his3Δ1 leu2Δ0*), and a S288C negative control strain (DMA1, *MAT****a*** *his3Δ::kanMX ura3Δ0 leu2Δ0 met15Δ0 or DMA809*, *MAT****a*** *hoΔ::kanMX his3Δ0 ura3Δ0 leu2Δ0 met15Δ0*; Data S8) were crossed to a collection of temperature sensitive mutants of essential genes (*MATα xxx-ts::natMX4 can1Δ::STE2pr-SpHIS5 lyp1Δ his3Δ1 leu2Δ0 ura3Δ0 met15Δ0*; (Costanzo et al. 2016)). SGA analysis was performed as described previously (Baryshnikova et al. 2010), with the exception that 5% mannose was added to the YPD plates used in the first steps of SGA analysis to facilitate pinning of the wild isolates. The final double mutant selection was performed at both 26 and 34 °C.

Plate images were processed with gitter v1.0.3 (Wagih and Parts 2014) and normalised with SGAtools (Wagih et al. 2013). Briefly, this process includes processing plate image files to detect the grid of colonies, quantifying the colony sizes, filtering out any technical replicates that accounted for at least 90% of the variation in the signal, averaging the remaining technical replicates, and log2-transforming. To calculate fitness values, we also averaged across multiple biological replicates. For the six out of ten wild strains that had two biological replicates, we used the replicate with the largest number of query strain measurements as a reference, and fit the other to it using linear regression. We used the single measurement for the remaining four strains. The average log2-scale colony size of all measurements passing the filters was reported as the fitness value at both permissive and restrictive temperatures. We filtered out 379 query strains that did not show lower fitness at the restrictive temperature (reference fitness difference between 26 and 34 °C below 0.2), retaining 1120 query strains in total for 580 genes. In addition, we removed strains for query genes that were genetically linked to the *HIS3*, *HO*, or *URA3* loci that were used for selection in the screen.

### SGA suppression analysis

To estimate suppression of the mutation effect by a wild strain, we quantified the difference in fitness at restrictive temperature after adjusting for overall growth between the reference and wild strains. To adjust, we set the median restrictive temperature fitnesses of temperature-insensitive strains to be equal, and scaled the wild strain restrictive temperature fitnesses to minimise mean-squared error of the fit to the respective values of reference. Importantly, growth at the permissive temperature, and its additional measurement noise, were not considered beyond filtering for strains that were not fit at the permissive temperature in the reference, as described above.

To generate genotype and phenotype trees, we used the scikit-learn average() function to compute the UPGMA tree, and the dendrogram() function for display (Pedregosa et al. 2012). For genotype trees, we calculated the distance between strains as the number of called genetic variants that are present in either strain, but not the other one. For phenotype trees, we calculated the distance between strains as 1 minus the Pearson’s correlation of their suppression profiles.

To test for consistency of suppression within complexes and pathways, we considered multiple functional annotation datasets. The sources for these datasets were: protein complexes (the Complex Portal (Meldal et al. 2015), downloaded June 6, 2018), KEGG pathway annotation (Kanehisa et al. 2016), coexpression degree (number of gene partners with a coexpression score > 1; (Huttenhower et al. 2006), and subcellular localisation (Huh et al. 2003). For each annotation that groups multiple genes, we calculated both complex average suppression value across TS alleles and wild strains, as well as average correlation of suppression values across wild strains between all possible allele pairs. To evaluate the significance of these values for a complex with *N* alleles, we sampled random alleles. For mean suppression, we sampled the *N* alleles 1,000 times, and computed the means. For mean correlation of suppression, we sampled the matching number of *N*(*N*-1)/2 allele pairs, while also matching the number of pairs that came from the same gene. We calculated the p-value of enrichment as the frequency of observing statistics from permuted data more extreme than the real value, and used the false discovery rate correction to adjust the p-values.

### Random sporulation assay

A total of 102 wild strain x TS allele combinations were selected for confirmation assays (Data S3). Between 3 to 20 different TS alleles were tested for each of the 10 wild strains, for a total of 78 different TS alleles corresponding to 56 different essential genes. The selected crosses spanned a wide range of suppression scores, and included 7 crosses with a negative suppression value. As controls, we crossed each selected TS allele to a reference S288C strain, and each wild strain was crossed to a wild-type S288C reference strain, giving a total of 78 S288C TS allele controls, and 10 S288C x wild strain controls.

All 190 strain pairs were crossed and sporulated. Sporulated cells were plated onto two agar plates that selected for haploid *MAT****a*** spores that carried the TS allele (SD −His/Arg/Lys +CAN/LYP/NAT). One plate was incubated at 26 °C and one at 34 °C. After 3 days plates were imaged, and colony size and number were determined using CellProfiler (Carpenter et al. 2006). We calculated the difference between the number of colonies and the colony area at 26 °C and 34 °C for each TS allele - wild strain combination, and compared the values for the S288C control to those of the wild strain crosses (Data S3). Images that contained <100 colonies at 26 °C were excluded from the analysis, and all images with <30 colonies were excluded from colony size determination. A TS allele - wild strain pair was considered to show suppression when either the number or the average size of the colonies of the wild cross was substantially larger than that of the control cross (TS allele x S288C) at 34 °C (Data S3).

### Sequencing, read mapping, SNP calling, and QTL analysis

We selected 38 crosses that showed various levels of suppression in the screen for bulk segregant analysis. The 38 samples included 6 positive controls involving query genes located on the left arm of chromosome XVI that were suppressed by the chrVIII-chrXVI translocation, 6 positive controls involving genes located on chromosome II or chromosome VIII that were suppressed by one of the NCYC110 aneuploidies, 6 cases that showed weak “suppression” in our screen (suppression score <0.6), and 20 cases that showed strong suppression in our screen (suppression score >0.75). In addition, we crossed each wild strain to a S288C reference strain. We collected at least two replicates of 1000 haploid progeny colonies per temperature for each cross, using the random sporulation assay outlined above. Colonies were scraped from the agar plates, and genomic DNA was isolated from the pools using the Qiagen DNeasy Blood & Tissue kit. Samples were sequenced using Illumina sequencing.

For each bulk segregant sequencing sample, we performed read mapping and variant calling under the Varathon framework (https://github.com/yjx1217/Varathon). Briefly, the raw reads were trimmed by trimmomatic v0.38 (Bolger, Lohse, and Usadel 2014) and subsequently mapped to the yeast reference genome (SGD R64-1-1) using bwa v0.7.17 (Li and Durbin 2009). The resulting read alignment was further processed by samtools v1.9 (Li et al. 2009), picard tools 2.18.25 (https://broadinstitute.github.io/picard/), and GATK3 v3.6 for sorting, duplicate removal, INDEL realignment and indexing. Variant calling was carried out by freebayes v1.2.0 (Garrison and Marth 2012) with the customised options “--ploidy 1 --min-alternate-fraction 0 --genotype-qualities”. Raw variant calls were processed by vt (github commit version f6d2b5d) (Tan, Abecasis, and Kang 2015) for variant decomposition, normalisation, annotation, and filtered by vcffilter (distributed together with freebayes) with the filter: “QUAL > 30 & QUAL / AO > 1 & SAF > 0 & SAR > 0 & RPR > 1 & RPL > 1”. Finally, VEP 101.0 (McLaren et al. 2016) was used to evaluate the functional impact of each variant by leveraging its specific genomic context.

We stratified the 38 bulk segregant QTL mapping experiments according to genomic coverage and screen signal. We separated the six crosses with the NCYC110 strain due to the wild strain ploidy issues, seven further crosses that had evidence for aneuploidy from sequencing coverage, and a final six crosses with chrXVI-VIII translocation that creates an additional wild type copy of the query gene in the segregants. To call QTLs in the remaining 19 samples without ploidy issues, and with strong or moderate suppression scores in the screen, we used Selection QTL Mapper (https://github.com/PMBio/sqtl), which implements the approach used for bulk segregant analysis mapping described in (Parts et al. 2011). Briefly, this approach first estimates reference allele frequencies in each sample using a probabilistic model that includes allele frequencies as latent variables, sequencing reads as observations, and the recombination rate parameter to couple frequencies at nearby sites. The posterior allele frequency distributions were then combined across biological replicates according to Bayes rule, and used to identify a broad set of QTL regions that had at least 12% frequency change between permissive and restrictive temperatures, and were at least 1kb long, using parameters “af_lenient=0.8, sd_lenient=3, af_stringent=0.12, sd_stringent=5, length_cutoff=1000, peak_cutoff=0.03”. A stricter set with allele frequency change of at least 0.20 was used for all but reproducibility analyses. Sites within 30kb of the TS allele or a SGA selection marker were not considered as QTL candidates.

All whole-genome sequencing data are publicly available at NCBI’s Sequence Read Archive (http://www.ncbi.nlm.nih.gov/sra), under accession number PRJNA673501.

### Suppressor gene prediction

For each detected QTL, we predicted the potential causal suppressor genes by ranking the genes for which the allele frequency change was within 3% of the strongest selected variant in the region by their functional relationship to the query gene, as described previously (van Leeuwen et al. 2020). In addition, we scored essential candidate genes higher than nonessential genes. Briefly, we evaluated the following functional relationships and gene properties in this order of priority: co-complex (highest priority), co-pathway, co-expression, co-localisation, and essentiality of the suppressor candidate (lowest priority). Thus, genes with co-complex relationships were ranked above those with only co-pathway relationships. Additionally, the order between genes within a given set was established by evaluating the rest of the functional relationships. For instance, the set of genes that were co-expressed with the query gene, but not in the same complex or pathway, were further ranked by whether they co-localised (highest rank) or not (lowest rank) with the query. The sources for these datasets were: protein complexes (the Complex Portal (Meldal et al. 2015), downloaded June 6, 2018), KEGG pathway annotation (Kanehisa et al. 2016), co-expression degree (number of gene partners with a co-expression score > 1; (Huttenhower et al. 2006), and subcellular localisation (Huh et al. 2003). We manually added suppressor candidate genes with genetic interactions or other known functional connections to the query gene that were not captured by our computational prediction, and also included known general modifier genes *MKT1* and *HAP1*.

### Suppressor candidate validation

To validate the predicted suppressor genes, we introduced 50 potential suppressor alleles into the reference genetic background. First, *kanR* or *nrsR* (“*natR*”) targeting guide RNA (gRNA) sequences were cloned into the pML104 or pML107 plasmid vectors, which carry Cas9 and either *URA3* or *LEU2* (Data S8, (Laughery et al. 2015)). Second, for nonessential suppressor gene candidates, we amplified the genes including ~400 bp upstream of the start codon and ~400 bp downstream of the stop codon from the various wild strains by PCR, and co-transformed the PCR fragment and the pML104-kanR1136 and pML107-kanR468 plasmids (Data S8) into a strain carrying a deletion allele of the suppressor gene (*MAT****a*** *xxxΔ::kanMX4 his3Δ1 leu2Δ0 ura3Δ0 met15Δ0*; Euroscarf). The gRNAs will cut the *kanMX4* cassette at two places, and the homology of the promoter and terminator sequences of the PCR product to the genomic sequences flanking the double-stranded DNA breaks will promote repair via homologous recombination and integration of the PCR product into the genome. For essential genes we used a similar strategy using a set of haploid strains in which the essential gene of interest was deleted in the genome but present on a plasmid (*MAT****a*** *xxxΔ::natR_kanR(Cterm) his3Δ1 leu2Δ0 ura3Δ0 +* [*XXX, URA3*]; (van Leeuwen et al. 2020)), and the plasmids pML104-natR412 and pML107-natR854 (Data S8) that carry gRNAs that target *natR*.

Transformants were initially selected on SD -Ura -Leu, and then propagated on YPD. Within 3 days of growth on YPD, the vast majority of yeast strains had lost the gRNA plasmids and properly replaced the suppressor candidate deletion allele with the wild allele, which we confirmed by PCR. For essential genes, we streaked the allele-swapped strains on SDall + 5-FOA to remove the plasmid carrying the essential gene.

Next, we crossed the allele-swapped strains to the corresponding TS mutant, and sporulated the resulting diploids. We isolated haploid progeny carrying the TS allele and confirmed the identity of the suppressor allele by Sanger sequencing. Growth of the TS mutants carrying the suppressor candidate allele from a wild strain was monitored at various temperatures to confirm the suppression phenotype.

Gain- or loss-of-function effects were predicted for each validated suppressor gene based on previously described genetic interactions between the query allele and deletion or overexpression alleles of the suppressor gene (Oughtred et al. 2019), or based on known phenotypes of the S288C and wild alleles (i.e. *HAP1* and *MKT1*).

### Smc6-FLAG chromatin immunoprecipitation

Smc6-FLAG strains were constructed by PCR gene-targeting (Longtine et al. 1998) using primers AGAGACCCTGAGAGACAGAATAATTCCAATTTTTATAATcggatccccgggttaattaa and GACGATTACACAATATTTTGAATAATTACATGAAGAAACAgcgcgttggccgattcatta to amplify the FLAG-tag from pFA6-6xGLY-3xFLAG-HIS3MX6 (Funakoshi and Hochstrasser 2009). Proper tagging was checked by colony PCR using primers TGCGGTCAAGGATTATTGCG and CGCTGTGAGAGTTGTTGAGG.

Smc6-FLAG expression was confirmed by western blotting. For each strain, whole cell extracts were prepared by TCA precipitation using 10 OD_600_-units of cells, and analyzed by SDS-PAGE. Western blotting was performed using an anti-FLAG antibody (clone M2, Sigma-Aldrich). Ponceau staining was used as a loading control.

Chromatin immunoprecipitation (ChIP) was performed as previously described with slight modifications (Cobb et al. 2003). Briefly, cells were grown to 5 x 10^6^ cells/ml in YPD and arrested in G2/M by incubation with nocodazole (15 μg/ml, Sigma-Aldrich) for 2 hours. Samples were fixed with 1 % formaldehyde. Cell pellets were resuspended in lysis buffer (50 mM Hepes, pH=7.5, 140 mM NaCl, 1 mM Na EDTA, 1% Triton X-100, 0.1 % sodium deoxycholate) containing protease inhibitors. Extracts were incubated with Dynabeads mouse IgG (Invitrogen, M-280) coated with antibody against FLAG (clone M2, Sigma-Aldrich) for 2 hours at 4°C. DNA was purified and enrichment at specific loci was measured using qPCR. Relative enrichment was determined by 2^−DDCt^ method (Livak and Schmittgen 2001; Cobb and van Attikum 2010). Dynabeads without antibody were used to correct for background. An amplicon 14 kb downstream of ARS607, devoid of Smc6 binding, was used for normalisation (Lindroos et al. 2006). Primers used are listed below.

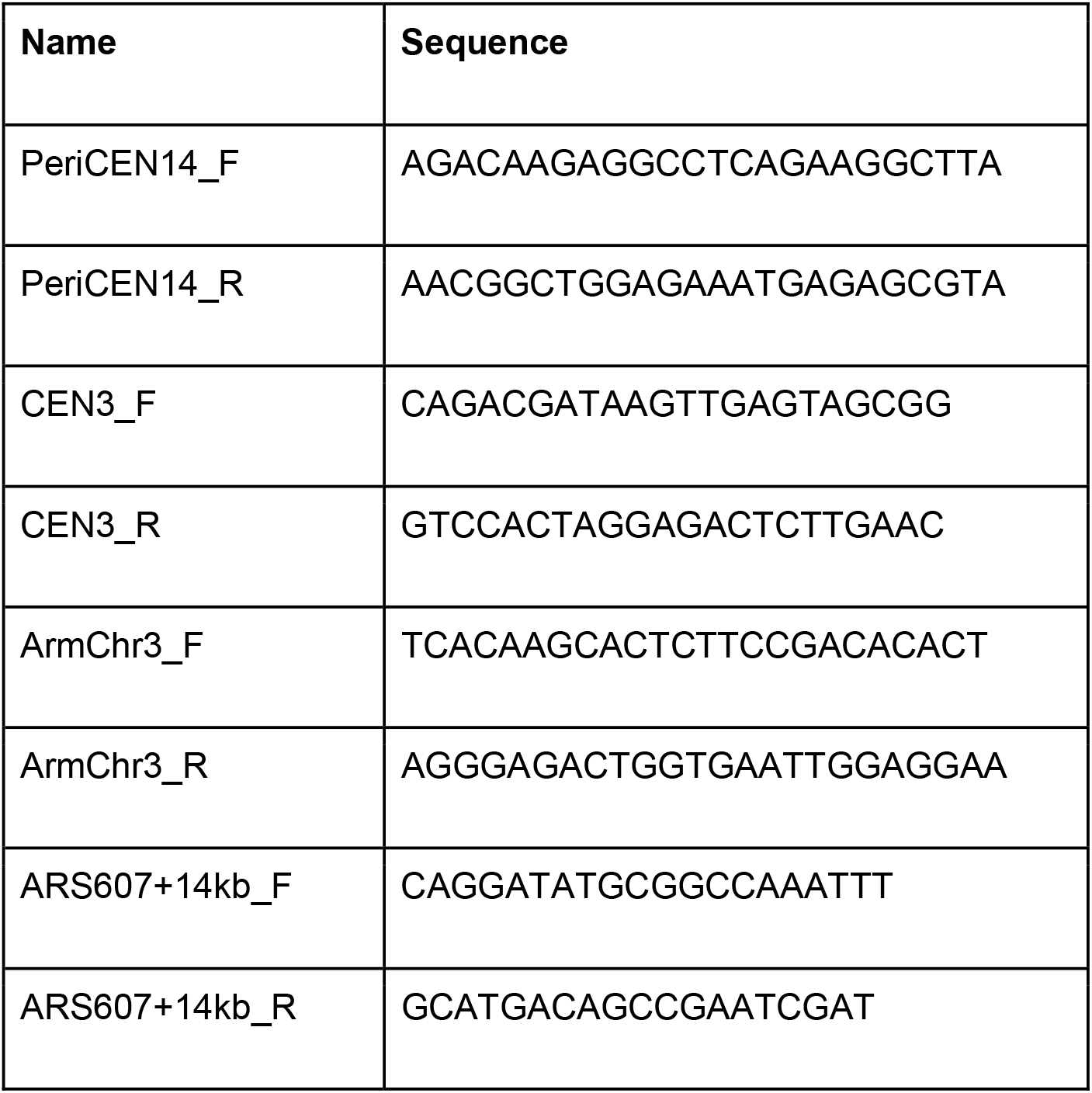

## References

Albert, Frank W., Joshua S. Bloom, Jake Siegel, Laura Day, and Leonid Kruglyak. 2018. “Genetics of Regulatory Variation in Gene Expression.” eLife 7 (July). https://doi.org/10.7554/eLife.35471.

Albert, Frank W., Sebastian Treusch, Arthur H. Shockley, Joshua S. Bloom, and Leonid Kruglyak. 2014. “Genetics of Single-Cell Protein Abundance Variation in Large Yeast Populations.” Nature 506 (7489): 494–97.

Altenburg, E., and H. J. Muller. 1920. “The Genetic Basis of Truncate Wing,-an Inconstant and Modifiable Character in Drosophila.” Genetics 5 (1): 1–59.

Bard, Jared A. M., Ellen A. Goodall, Eric R. Greene, Erik Jonsson, Ken C. Dong, and Andreas Martin. 2018. “Structure and Function of the 26S Proteasome.” Annual Review of Biochemistry 87 (June): 697–724.

Barson, Nicola J., Tutku Aykanat, Kjetil Hindar, Matthew Baranski, Geir H. Bolstad, Peder Fiske, Céleste Jacq, et al. 2015. “Sex-Dependent Dominance at a Single Locus Maintains Variation in Age at Maturity in Salmon.” Nature 528 (7582): 405–8.

Baryshnikova, Anastasia, Michael Costanzo, Yungil Kim, Huiming Ding, Judice Koh, Kiana Toufighi, Ji-Young Youn, et al. 2010. “Quantitative Analysis of Fitness and Genetic Interactions in Yeast on a Genome Scale.” Nature Methods 7 (12): 1017–24.

Behan, Fiona M., Francesco Iorio, Gabriele Picco, Emanuel Gonçalves, Charlotte M. Beaver, Giorgia Migliardi, Rita Santos, et al. 2019. “Prioritization of Cancer Therapeutic Targets Using CRISPR-Cas9 Screens.” Nature 568 (7753): 511–16.

Bergström, Anders, Jared T. Simpson, Francisco Salinas, Benjamin Barré, Leopold Parts, Amin Zia, Alex N. Nguyen Ba, et al. 2014. “A High-Definition View of Functional Genetic Variation from Natural Yeast Genomes.” Molecular Biology and Evolution 31 (4): 872–88.

Bloom, Joshua S., James Boocock, Sebastian Treusch, Meru J. Sadhu, Laura Day, Holly Oates-Barker, and Leonid Kruglyak. 2019. “Rare Variants Contribute Disproportionately to Quantitative Trait Variation in Yeast.” eLife 8 (October). https://doi.org/10.7554/eLife.49212.

Bolger, Anthony M., Marc Lohse, and Bjoern Usadel. 2014. “Trimmomatic: A Flexible Trimmer for Illumina Sequence Data.” Bioinformatics 30 (15): 2114–20.

Carpenter, Anne E., Thouis R. Jones, Michael R. Lamprecht, Colin Clarke, In Han Kang, Ola Friman, David A. Guertin, et al. 2006. “CellProfiler: Image Analysis Software for Identifying and Quantifying Cell Phenotypes.” Genome Biology 7 (10): R100.

Chandler, Christopher H., Sudarshan Chari, David Tack, and Ian Dworkin. 2014. “Causes and Consequences of Genetic Background Effects Illuminated by Integrative Genomic Analysis.” Genetics 196 (4): 1321–36.

Cobb, Jennifer A., Lotte Bjergbaek, Kenji Shimada, Christian Frei, and Susan M. Gasser. 2003. “DNA Polymerase Stabilization at Stalled Replication Forks Requires Mec1 and the RecQ Helicase Sgs1.” The EMBO Journal 22 (16): 4325–36.

Cobb, Jennifer, and Haico van Attikum. 2010. “Mapping Genomic Targets of DNA Helicases by Chromatin Immunoprecipitation in Saccharomyces Cerevisiae.” Methods in Molecular Biology 587: 113–26.

Costanzo, Michael, Benjamin VanderSluis, Elizabeth N. Koch, Anastasia Baryshnikova, Carles Pons, Guihong Tan, Wen Wang, et al. 2016. “A Global Genetic Interaction Network Maps a Wiring Diagram of Cellular Function.” Science 353 (6306). https://doi.org/10.1126/science.aaf1420.

Cubillos, Francisco A., Edward J. Louis, and Gianni Liti. 2009. “Generation of a Large Set of Genetically Tractable Haploid and Diploid Saccharomyces Strains.” FEMS Yeast Research 9 (8): 1217–25.

De Piccoli, Giacomo, Felipe Cortes-Ledesma, Gregory Ira, Jordi Torres-Rosell, Stefan Uhle, Sarah Farmer, Ji-Young Hwang, et al. 2006. “Smc5-Smc6 Mediate DNA Double-Strand-Break Repair by Promoting Sister-Chromatid Recombination.” Nature Cell Biology 8 (9): 1032–34.

Dowell, Robin D., Owen Ryan, An Jansen, Doris Cheung, Sudeep Agarwala, Timothy Danford, Douglas A. Bernstein, et al. 2010. “Genotype to Phenotype: A Complex Problem.” Science 328 (5977): 469.

Fay, Justin C. 2013. “The Molecular Basis of Phenotypic Variation in Yeast.” Current Opinion in Genetics & Development 23 (6): 672–77.

Funakoshi, Minoru, and Mark Hochstrasser. 2009. “Small Epitope-Linker Modules for PCR-Based C-Terminal Tagging in Saccharomyces Cerevisiae.” Yeast 26 (3): 185–92.

Galardini, Marco, Bede P. Busby, Cristina Vieitez, Alistair S. Dunham, Athanasios Typas, and Pedro Beltrao. 2019. “The Impact of the Genetic Background on Gene Deletion Phenotypes in Saccharomyces Cerevisiae.” Molecular Systems Biology 15 (12): e8831.

Garrison, Erik, and Gabor Marth. 2012. “Haplotype-Based Variant Detection from Short-Read Sequencing.” http://arxiv.org/abs/1207.3907.

Giaever, Guri, Angela M. Chu, Li Ni, Carla Connelly, Linda Riles, Steeve Véronneau, Sally Dow, et al. 2002. “Functional Profiling of the Saccharomyces Cerevisiae Genome.” Nature 418 (6896): 387–91.

Gonçalves, Emanuel, Aldo Segura-Cabrera, Clare Pacini, Gabriele Picco, Fiona M. Behan, Patricia Jaaks, Elizabeth A. Coker, et al. 2020. “Drug Mechanism-of-Action Discovery through the Integration of Pharmacological and CRISPR Screens.” Molecular Systems Biology 16 (7): e9405.

Hallin, Johan, Kaspar Martens, Alexander Young, Martin Zackrisson, Francisco Salinas, Leopold Parts, Jonas Warringer, and Gianni Liti. 2016. “Powerful Decomposition of Complex Traits in a Diploid Model Using Phased Outbred Lines.” Nature Communications 7:1311.

Hamilton, Bruce A., and Benjamin D. Yu. 2012. “Modifier Genes and the Plasticity of Genetic Networks in Mice.” PLoS Genetics 8 (4): e1002644.

Harper, Andrew R., Shalini Nayee, and Eric J. Topol. 2015. “Protective Alleles and Modifier Variants in Human Health and Disease.” Nature Reviews. Genetics 16 (12): 689–701.

Hose, James, Chris Mun Yong, Maria Sardi, Zhishi Wang, Michael A. Newton, and Audrey P. Gasch. 2015. “Dosage Compensation Can Buffer Copy-Number Variation in Wild Yeast.” eLife 4 (May). https://doi.org/10.7554/eLife.05462.

Hou, Jing, Jolanda van Leeuwen, Brenda J. Andrews, and Charles Boone. 2018. “Genetic Network Complexity Shapes Background-Dependent Phenotypic Expression.” Trends in Genetics: TIG 34 (8): 578–86.

Hou, Jing, Guihong Tan, Gerald R. Fink, Brenda J. Andrews, and Charles Boone. 2019. “Complex Modifier Landscape Underlying Genetic Background Effects.” Proceedings of the National Academy of Sciences of the United States of America 116 (11): 5045–54.

Hudson, Jessica J. R., Katerina Bednarova, Lucie Kozakova, Chunyan Liao, Marc Guerineau, Rita Colnaghi, Susanne Vidot, et al. 2011. “Interactions between the Nse3 and Nse4 Components of the SMC5-6 Complex Identify Evolutionarily Conserved Interactions between MAGE and EID Families.” PloS One 6 (2): e17270.

Huh, Won-Ki, James V. Falvo, Luke C. Gerke, Adam S. Carroll, Russell W. Howson, Jonathan S. Weissman, and Erin K. O’Shea. 2003. “Global Analysis of Protein Localization in Budding Yeast.” Nature 425 (6959): 686–91.

Huttenhower, C., M. Hibbs, C. Myers, and O. G. Troyanskaya. 2006. “A Scalable Method for Integration and Functional Analysis of Multiple Microarray Datasets.” Bioinformatics. https://doi.org/10.1093/bioinformatics/btl492.

Jeppsson, Kristian, Kristian K. Carlborg, Ryuichiro Nakato, Davide G. Berta, Ingrid Lilienthal, Takaharu Kanno, Arne Lindqvist, et al. 2014. “The Chromosomal Association of the Smc5/6 Complex Depends on Cohesion and Predicts the Level of Sister Chromatid Entanglement.” PLoS Genetics 10 (10): e1004680.

Johnston, Susan E., John C. McEwan, Natalie K. Pickering, James W. Kijas, Dario Beraldi, Jill G. Pilkington, Josephine M. Pemberton, and Jon Slate. 2011. “Genome-Wide Association Mapping Identifies the Genetic Basis of Discrete and Quantitative Variation in Sexual Weaponry in a Wild Sheep Population.” Molecular Ecology 20 (12): 2555–66.

Jones, Matthew R., L. Scott Mills, Paulo Célio Alves, Colin M. Callahan, Joel M. Alves, Diana J. R. Lafferty, Francis M. Jiggins, Jeffrey D. Jensen, José Melo-Ferreira, and Jeffrey M. Good. 2018. “Adaptive Introgression Underlies Polymorphic Seasonal Camouflage in Snowshoe Hares.” Science 360 (6395): 1355–58.

Kanehisa, Minoru, Yoko Sato, Masayuki Kawashima, Miho Furumichi, and Mao Tanabe. 2016. “KEGG as a Reference Resource for Gene and Protein Annotation.” Nucleic Acids Research 44 (D1): D457–62.

Laughery, Marian F., Tierra Hunter, Alexander Brown, James Hoopes, Travis Ostbye, Taven Shumaker, and John J. Wyrick. 2015. “New Vectors for Simple and Streamlined CRISPR-Cas9 Genome Editing in Saccharomyces Cerevisiae.” Yeast 32 (12): 711–20.

Leeuwen, Jolanda van, Carles Pons, Charles Boone, and Brenda J. Andrews. 2017. “Mechanisms of Suppression: The Wiring of Genetic Resilience.” BioEssays: News and Reviews in Molecular, Cellular and Developmental Biology 39 (7). https://doi.org/10.1002/bies.201700042.

Leeuwen, Jolanda van, Carles Pons, Joseph C. Mellor, Takafumi N. Yamaguchi, Helena Friesen, John Koschwanez, Mojca Mattiazzi Ušaj, et al. 2016. “Exploring Genetic Suppression Interactions on a Global Scale.” Science 354 (6312). https://doi.org/10.1126/science.aag0839.

Leeuwen, Jolanda van, Carles Pons, Guihong Tan, Jason Zi Wang, Jing Hou, Jochen Weile, Marinella Gebbia, et al. 2020. “Systematic Analysis of Bypass Suppression of Essential Genes.” Molecular Systems Biology 16 (9): e9828.

Li, Heng, and Richard Durbin. 2009. “Fast and Accurate Short Read Alignment with Burrows-Wheeler Transform.” Bioinformatics 25 (14): 1754–60.

Li, H., B. Handsaker, A. Wysoker, T. Fennell, J. Ruan, N. Homer, G. Marth, G. Abecasis, R. Durbin, and 1000 Genome Project Data Processing Subgroup. 2009. “The Sequence Alignment/Map Format and SAMtools.” Bioinformatics. https://doi.org/10.1093/bioinformatics/btp352.

Li, Jing, Ignacio Vázquez-García, Karl Persson, Asier González, Jia-Xing Yue, Benjamin Barré, Michael N. Hall, et al. 2019. “Shared Molecular Targets Confer Resistance over Short and Long Evolutionary Timescales.” Molecular Biology and Evolution 36 (4): 691–708.

Lindroos, Hanna Betts, Lena Ström, Takehiko Itoh, Yuki Katou, Katsuhiko Shirahige, and Camilla Sjögren. 2006. “Chromosomal Association of the Smc5/6 Complex Reveals That It Functions in Differently Regulated Pathways.” Molecular Cell 22 (6): 755–67.

Liti, Gianni, David M. Carter, Alan M. Moses, Jonas Warringer, Leopold Parts, Stephen A. James, Robert P. Davey, et al. 2009. “Population Genomics of Domestic and Wild Yeasts.” Nature 458 (7236): 337–41.

Liti, Gianni, and Edward J. Louis. 2012. “Advances in Quantitative Trait Analysis in Yeast.” PLoS Genetics 8 (8): e1002912.

Livak, K. J., and T. D. Schmittgen. 2001. “Analysis of Relative Gene Expression Data Using Real-Time Quantitative PCR and the 2(-Delta Delta C(T)) Method.” Methods 25 (4): 402–8.

Longtine, M. S., A. McKenzie 3rd, D. J. Demarini, N. G. Shah, A. Wach, A. Brachat, P. Philippsen, and J. R. Pringle. 1998. “Additional Modules for Versatile and Economical PCR-Based Gene Deletion and Modification in Saccharomyces Cerevisiae.” Yeast 14 (10): 953–61.

Magtanong, Leslie, Cheuk Hei Ho, Sarah L. Barker, Wei Jiao, Anastasia Baryshnikova, Sondra Bahr, Andrew M. Smith, et al. 2011. “Dosage Suppression Genetic Interaction Networks Enhance Functional Wiring Diagrams of the Cell.” Nature Biotechnology 29 (6): 505–11.

Matsui, Takeshi, Jonathan T. Lee, and Ian M. Ehrenreich. 2017. “Genetic Suppression: Extending Our Knowledge from Lab Experiments to Natural Populations.” BioEssays: News and Reviews in Molecular, Cellular and Developmental Biology 39 (7). https://doi.org/10.1002/bies.201700023.

McLaren, William, Laurent Gil, Sarah E. Hunt, Harpreet Singh Riat, Graham R. S. Ritchie, Anja Thormann, Paul Flicek, and Fiona Cunningham. 2016. “The Ensembl Variant Effect Predictor.” Genome Biology. https://doi.org/10.1186/s13059-016-0974-4.

Meldal, Birgit H. M., Oscar Forner-Martinez, Maria C. Costanzo, Jose Dana, Janos Demeter, Marine Dumousseau, Selina S. Dwight, et al. 2015. “The Complex Portal--an Encyclopaedia of Macromolecular Complexes.” Nucleic Acids Research 43 (Database issue): D479–84.

Menolfi, Demis, Axel Delamarre, Armelle Lengronne, Philippe Pasero, and Dana Branzei. 2015. “Essential Roles of the Smc5/6 Complex in Replication through Natural Pausing Sites and Endogenous DNA Damage Tolerance.” Molecular Cell 60 (6): 835–46.

Oughtred, Rose, Chris Stark, Bobby-Joe Breitkreutz, Jennifer Rust, Lorrie Boucher, Christie Chang, Nadine Kolas, et al. 2019. “The BioGRID Interaction Database: 2019 Update.” Nucleic Acids Research 47 (D1): D529–41.

Parts, Leopold, Francisco A. Cubillos, Jonas Warringer, Kanika Jain, Francisco Salinas, Suzannah J. Bumpstead, Mikael Molin, et al. 2011. “Revealing the Genetic Structure of a Trait by Sequencing a Population under Selection.” Genome Research 21 (7): 1131–38.

Parts, Leopold, Yi-Chun Liu, Manu M. Tekkedil, Lars M. Steinmetz, Amy A. Caudy, Andrew G. Fraser, Charles Boone, Brenda J. Andrews, and Adam P. Rosebrock. 2014. “Heritability and Genetic Basis of Protein Level Variation in an Outbred Population.” Genome Research 24 (8): 1363–70.

Pebernard, Stephanie, J. Jefferson P. Perry, John A. Tainer, and Michael N. Boddy. 2008. “Nse1 RING-like Domain Supports Functions of the Smc5-Smc6 Holocomplex in Genome Stability.” Molecular Biology of the Cell 19 (10): 4099–4109.

Pedregosa, Fabian, Gaël Varoquaux, Alexandre Gramfort, Vincent Michel, Bertrand Thirion, Olivier Grisel, Mathieu Blondel, et al. 2012. “Scikit-Learn: Machine Learning in Python.” http://arxiv.org/abs/1201.0490.

Pérez-Ortín, José E., Amparo Querol, Sergi Puig, and Eladio Barrio. 2002. “Molecular Characterization of a Chromosomal Rearrangement Involved in the Adaptive Evolution of Yeast Strains.” Genome Research 12 (10): 1533–39.

Peter, Jackson, Matteo De Chiara, Anne Friedrich, Jia-Xing Yue, David Pflieger, Anders Bergström, Anastasie Sigwalt, et al. 2018. “Genome Evolution across 1,011 Saccharomyces Cerevisiae Isolates.” Nature 556 (7701): 339–44.

Riordan, Jesse D., and Joseph H. Nadeau. 2017. “From Peas to Disease: Modifier Genes, Network Resilience, and the Genetics of Health.” American Journal of Human Genetics 101 (2): 177–91.

Sanchez, Monica R., Celia Payen, Frances Cheong, Blake T. Hovde, Sarah Bissonnette, Adam P. Arkin, Jeffrey M. Skerker, Rachel B. Brem, Amy A. Caudy, and Maitreya J. Dunham. 2019. “Transposon Insertional Mutagenesis in Saccharomyces Uvarum Reveals Trans-Acting Effects Influencing Species-Dependent Essential Genes.” Genome Research. https://doi.org/10.1101/gr.232330.117.

Steinmetz, Lars M., Himanshu Sinha, Dan R. Richards, Jamie I. Spiegelman, Peter J. Oefner, John H. McCusker, and Ronald W. Davis. 2002. “Dissecting the Architecture of a Quantitative Trait Locus in Yeast.” Nature 416 (6878): 326–30.

Tan, Adrian, Gonçalo R. Abecasis, and Hyun Min Kang. 2015. “Unified Representation of Genetic Variants.” Bioinformatics. https://doi.org/10.1093/bioinformatics/btv112.

Taylor, Matthew B., and Ian M. Ehrenreich. 2015. “Higher-Order Genetic Interactions and Their Contribution to Complex Traits.” Trends in Genetics: TIG 31 (1): 34–40.

Thompson, Neil F., Eric C. Anderson, Anthony J. Clemento, Matthew A. Campbell, Devon E. Pearse, James W. Hearsey, Andrew P. Kinziger, and John Carlos Garza. 2020. “A Complex Phenotype in Salmon Controlled by a Simple Change in Migratory Timing.” Science 370 (6516): 609–13.

Tong, A. H., M. Evangelista, A. B. Parsons, H. Xu, G. D. Bader, N. Pagé, M. Robinson, et al. 2001. “Systematic Genetic Analysis with Ordered Arrays of Yeast Deletion Mutants.” Science 294 (5550): 2364–68.

Wagih, Omar, and Leopold Parts. 2014. “Gitter: A Robust and Accurate Method for Quantification of Colony Sizes from Plate Images.” G3 4 (3): 547–52.

Wagih, Omar, Matej Usaj, Anastasia Baryshnikova, Benjamin VanderSluis, Elena Kuzmin, Michael Costanzo, Chad L. Myers, Brenda J. Andrews, Charles M. Boone, and Leopold Parts. 2013. “SGAtools: One-Stop Analysis and Visualization of Array-Based Genetic Interaction Screens.” Nucleic Acids Research 41 (Web Server issue): W591–96.

Wright, Caroline F., Ben West, Marcus Tuke, Samuel E. Jones, Kashyap Patel, Thomas W. Laver, R. N. Beaumont, et al. 2019. “Assessing the Pathogenicity, Penetrance and Expressivity of Putative Disease-Causing Variants in a Population Setting.” American Journal of Human Genetics 104 (2): 275–286.

