## Supplementary material for "Diverse natural variants suppress mutations in hundreds of essential genes": Figure S1

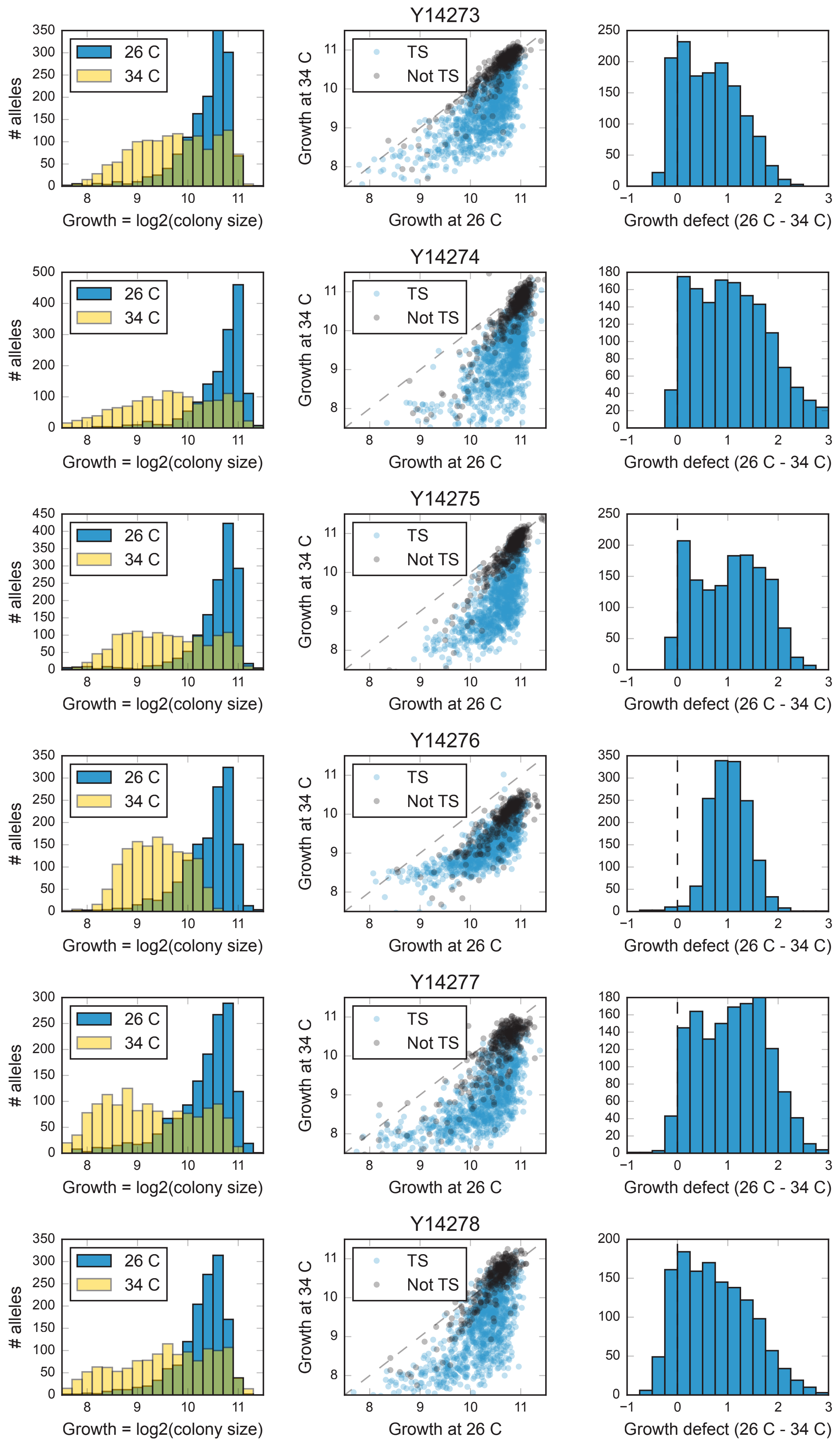


Continued on the next page

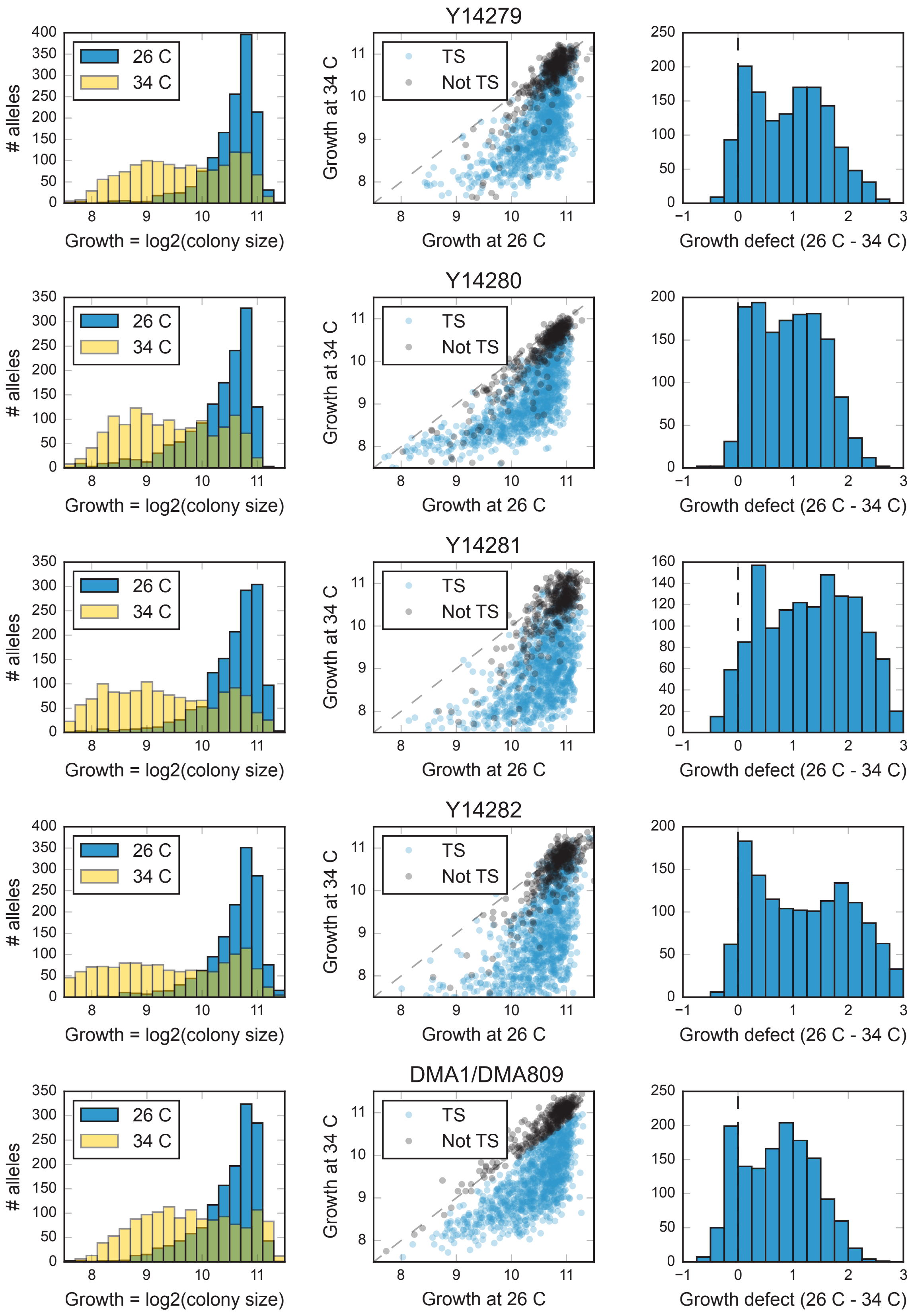


**Fig. S1. Distribution of fitness values at 26 and 34 °C.** Rows: ten wild strains and reference. Left column: frequency (y-axis) of TS allele to wild strain cross progeny growth (x-axis; log2-scaled colony size in pixels) at 26 °C (blue) and 34 °C (yellow). Middle column: growth at 26 °C (x-axis) and 34 °C (y-axis) of the same progeny as in left column (markers). Black: TS alleles deemed not temperature sensitive based on the control cross. Blue: all other TS alleles. Right column: frequency (y-axis) of growth defect (growth at 26 °C minus growth at 34 °C; x-axis). Y142.. and DMA.. labels correspond to the various wild and reference strains, respectively (see Data S8).
