## Supplementary material for "Diverse natural variants suppress mutations in hundreds of essential genes": Figure S2

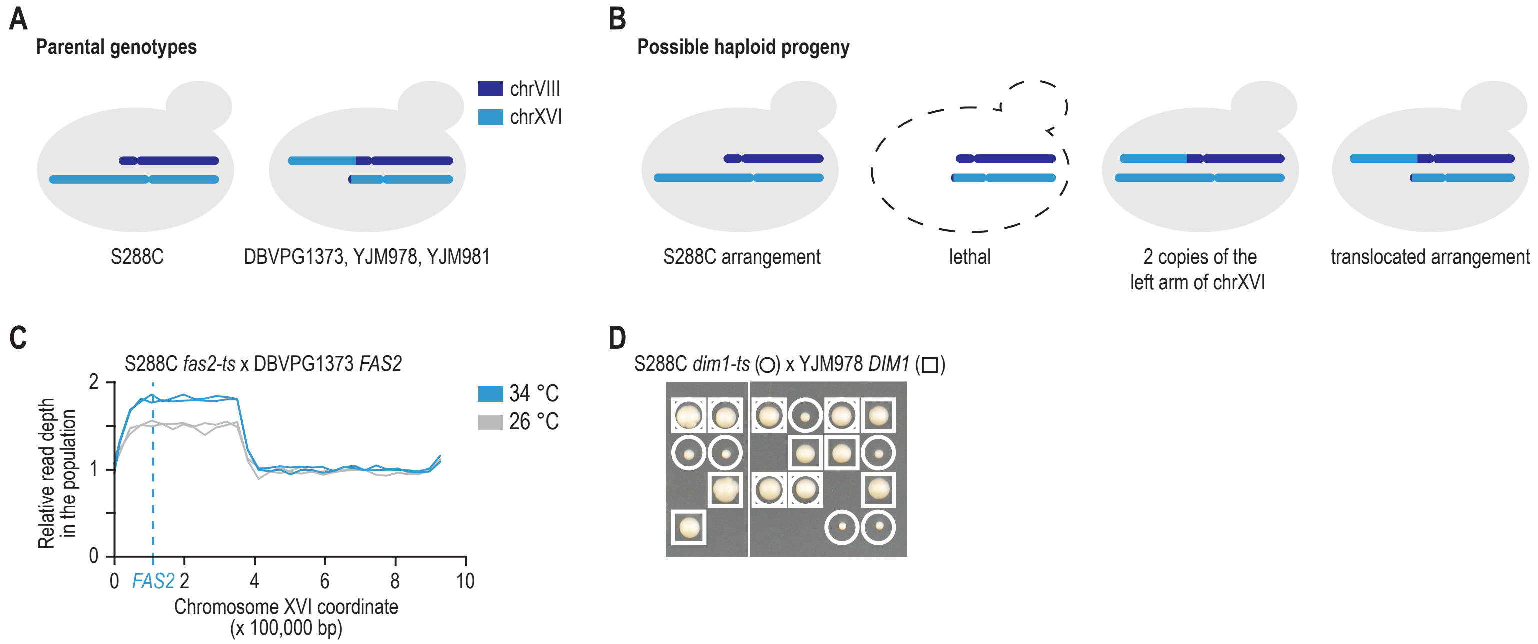


**Fig. S2. Suppression by a translocation between chrVIII and chrXVI.** (**A**) Three of the strains used in our screens carried a translocation between the promoter regions of *ECM34*, located at the left arm of chrVIII, and *SSU1*, located at the left arm of chrXVI (Pérez-Ortín et al., 2002). (**B**) Crossing S228C to one of the strains carrying the translocation will result in 25% spore lethality, due to the absence of a large part of the left arm of chrXVI, which is essential for viability. Of the viable spores, one third will carry the S288C chromosome arrangement, one third will carry the arrangement of the translocated strain, and one third will carry the S288C version of chrXVI, combined with the rearranged chromosome that carries the chrVIII centromere. This last combination of chromosomes will lead to the presence of two copies of a large part of the left arm of chrXVI. (**C**-**D**) A second copy of part of the left arm of chrXVI can suppress the TS phenotype of TS alleles located on this part of chrXVI. *FAS2* (*YPL231W*) and *DIM1* (*YPL266W*) are located on the translocated left arm of chrXVI. (C) Population sequencing read depth of haploid progeny carrying a *fas2-ts* allele, isolated from a cross between a S228c *fas2-ts* mutant and a DBVPG1373 wild-type strain, at either 26 or 34 °C. Part of the left arm of chrXVI is present at increased copy number in the population at 26 °C, due to the lethality of spores lacking the chromosome arm. At 34 °C, selection occurs for cells carrying two copies of this chromosomal fragment, because in addition to carrying the TS allele, they will also carry a wild-type copy of *FAS2* that is inherited from DBVPG1373. (D) Tetrad dissection of a cross between a S288C *dim1-ts* mutant and a YJM978 wild-type strain. Cells carrying both a *dim1-ts* and a *DIM1* wild-type allele have increased fitness compared to mutants carrying only a *dim1-ts* allele.
