## Supplementary material for "Diverse natural variants suppress mutations in hundreds of essential genes": Figure S3

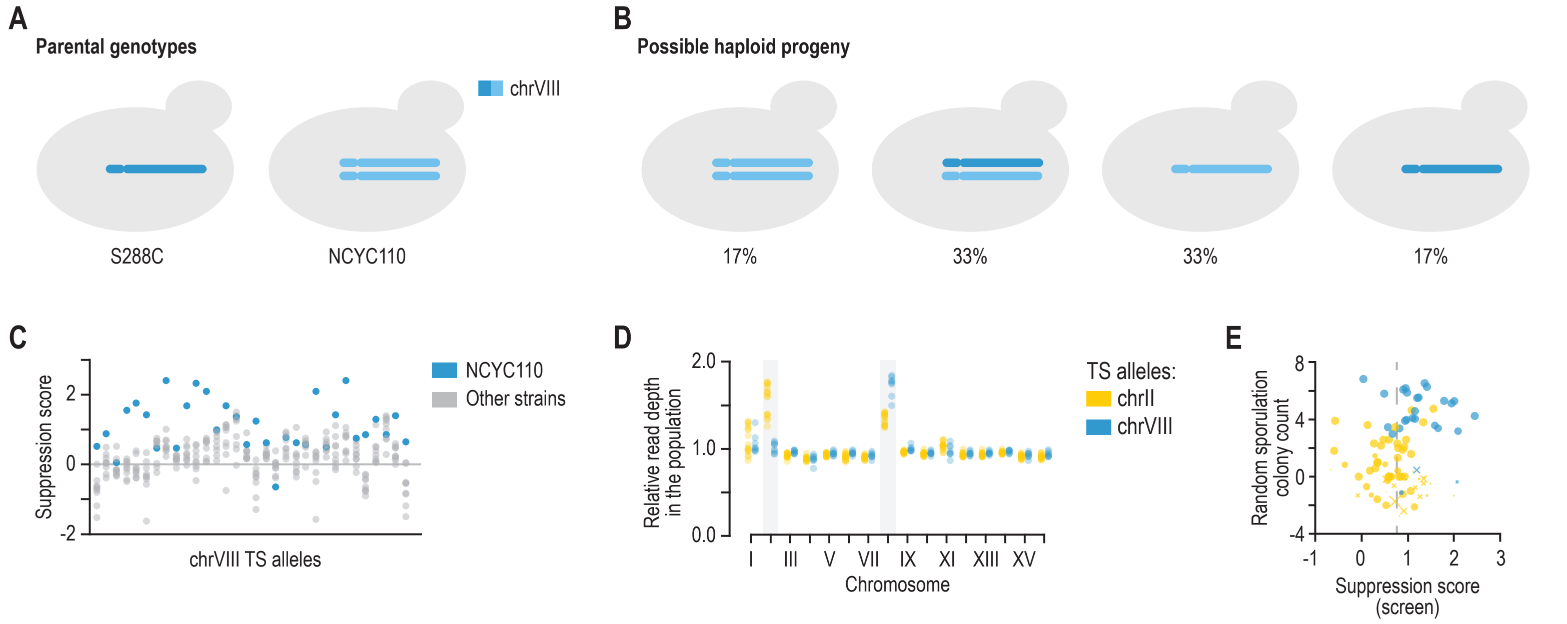


**Fig. S3. Aneuploidies and suppression validation.** (**A-B**) The NCYC110 strain that was used in our screen carried aneuploidies of chromosomes II and VIII. Chromosome VIII is highlighted as an example. Crossing S228C to NCYC110 will result in 50% of the progeny carrying one copy of chrVIII, and 50% of the progeny carrying two copies of chrVIII. Of the progeny carrying one copy of chrVIII, one third will carry the S288C chromosome and two thirds the NCYC110 chromosome. Of the progeny carrying two copies of chrVIII, one third will carry two NCYC110 chromosomes, and one third will carry one S288C and one NCYC110 version of chrVIII. We note that recombination can occur between the S288C and NCYC110 chromosomes. (**C-D**) A second copy of a chromosome can suppress the TS phenotype of TS alleles located on this chromosome. (C) For each TS allele located on chrVIII, the suppression scores are plotted for haploid progeny of a cross to NCYC110 (blue) or other wild isolates (grey). (D) Shown are the relative sequencing read depth of populations of haploid progeny isolated from a cross of NCYC110 to TS alleles located on either chrII (yellow) or chrVIII (blue). Cells carrying a TS allele on chrVIII have substantially higher coverage of this chromosome compared to query genes located on chrII. Note that aneuploidy of chromosome II is lost in the absence of positive selection, suggesting that it is detrimental in the population. (**E**) Screen and followup scores are concordant. Screen suppression score (x-axis; log2-scale growth ratio between wild strain and reference crosses at 34 °C; Methods) and individual followup colony count difference (y-axis; log2-scale colony count ratio in the wild strain cross between 26 and 34 °C minus log2-scale colony count ratio in the reference cross between 26 and 34 °C). Marker size: followup experiment colony count at permissive temperature (smaller markers for fewer colonies); marker type: “x” for TS strains with low temperature sensitivity in the followup control cross (colony count ratio between 26 and 34 °C below 2); “o” for the rest.
