## Supplementary material for "Diverse natural variants suppress mutations in hundreds of essential genes": Figure S4

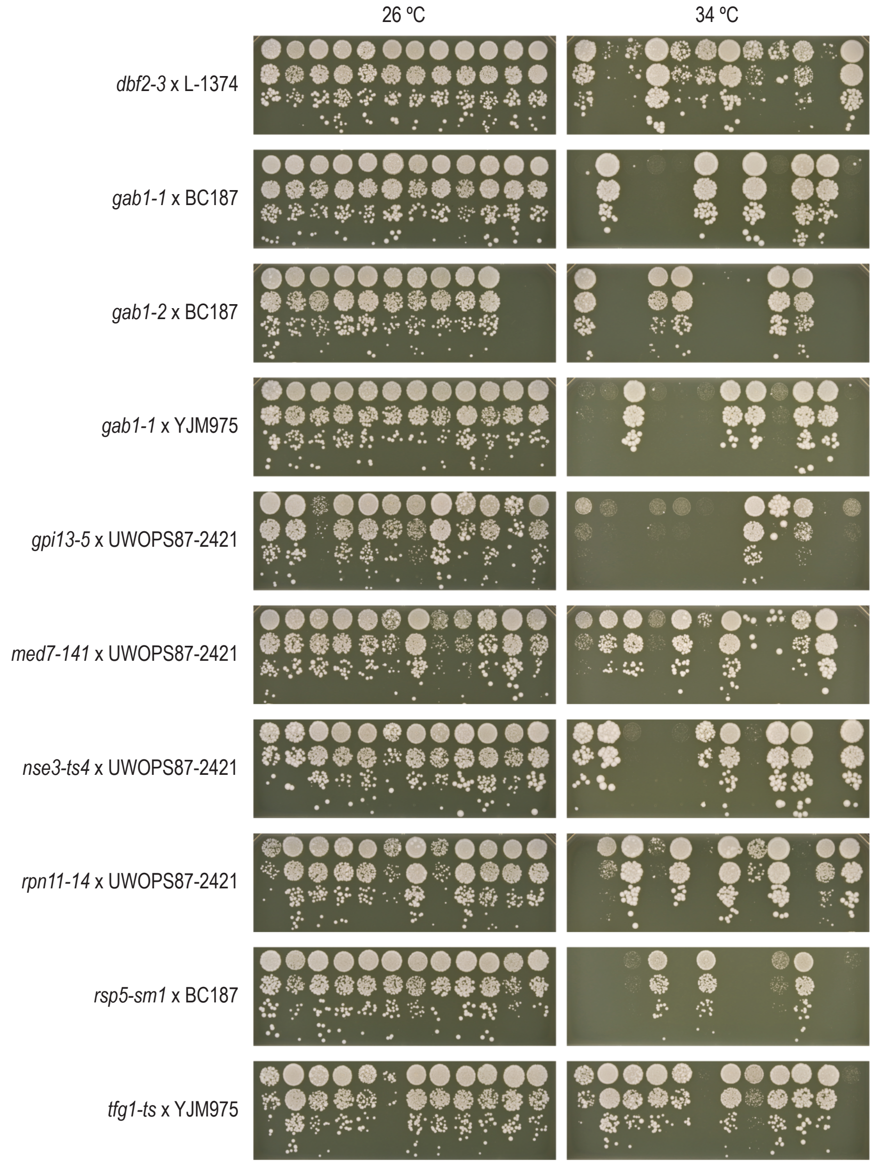


**Fig. S4. Estimating the number of modifiers.** TS alleles were crossed to the wild strain in which they were suppressed, and the resulting hybrid strains were dissected at 26 ºC. Twelve spores carrying the TS allele were randomly selected from the dissection plate, and were grown overnight in liquid media. Cultures were diluted to an optical density at 600 nm of 0.1 and a series of ten-fold dilutions was spotted on agar plates and incubated for 2 days at 26 ºC or 34 ºC. The number of modifiers was estimated by determining the fraction of the spores that grew well at 34 ºC.
