## Supplementary material for "Diverse natural variants suppress mutations in hundreds of essential genes": Figure S5

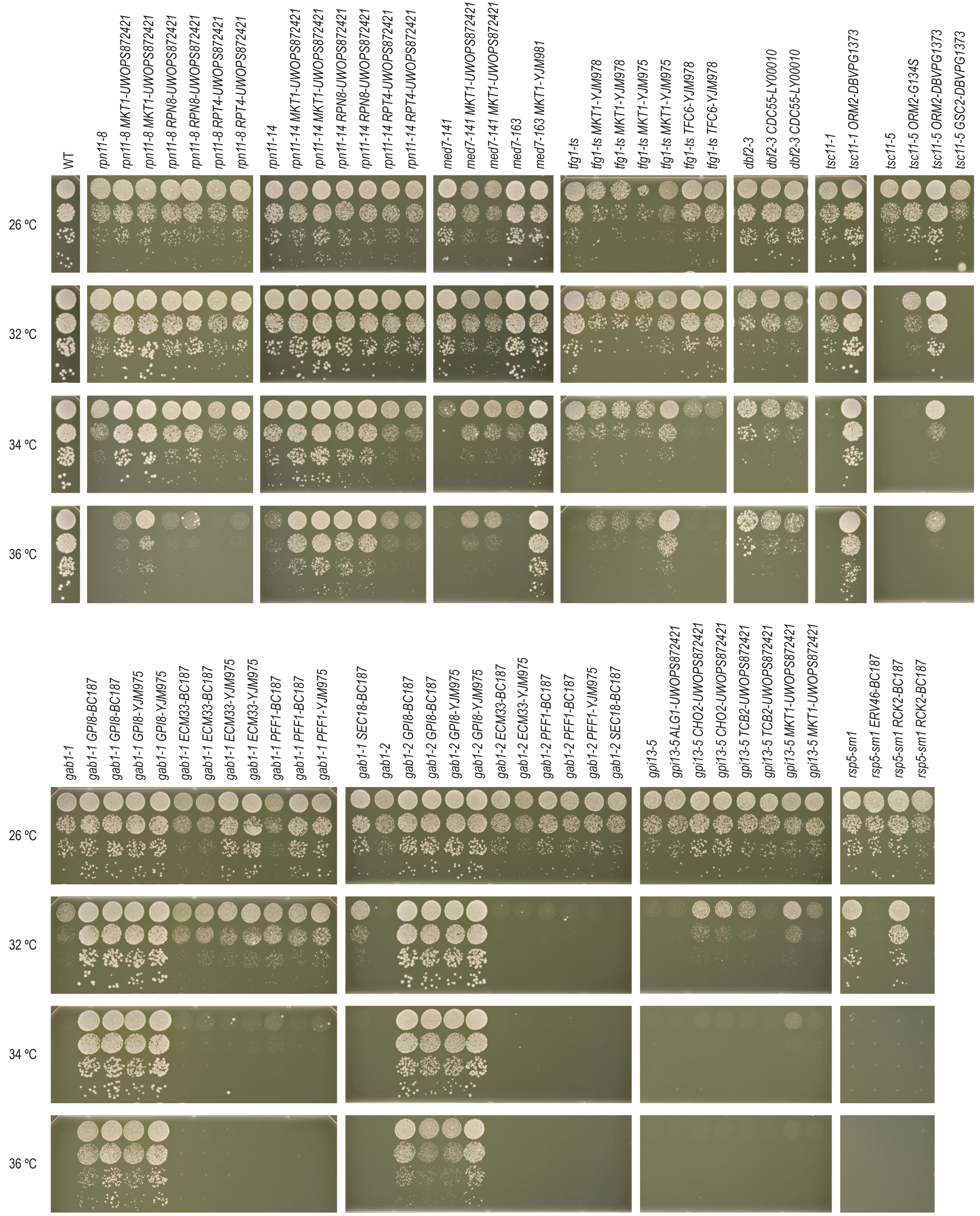


**Fig. S5. Suppressor identification.** To validate the suppressor candidates, we replaced the suppressor gene reference alleles with their wild version in the reference strain background, and tested for suppression of the corresponding TS allele. Cultures of the indicated strains were grown until saturation, and a series of ten-fold dilutions was spotted on YPD agar plates and incubated at the indicated temperatures. See Figure 5A for the suppression of *nse3-ts4* by *NSE1-UWOPS872421.*
