## Supplementary material for "Diverse natural variants suppress mutations in hundreds of essential genes": Figure S6

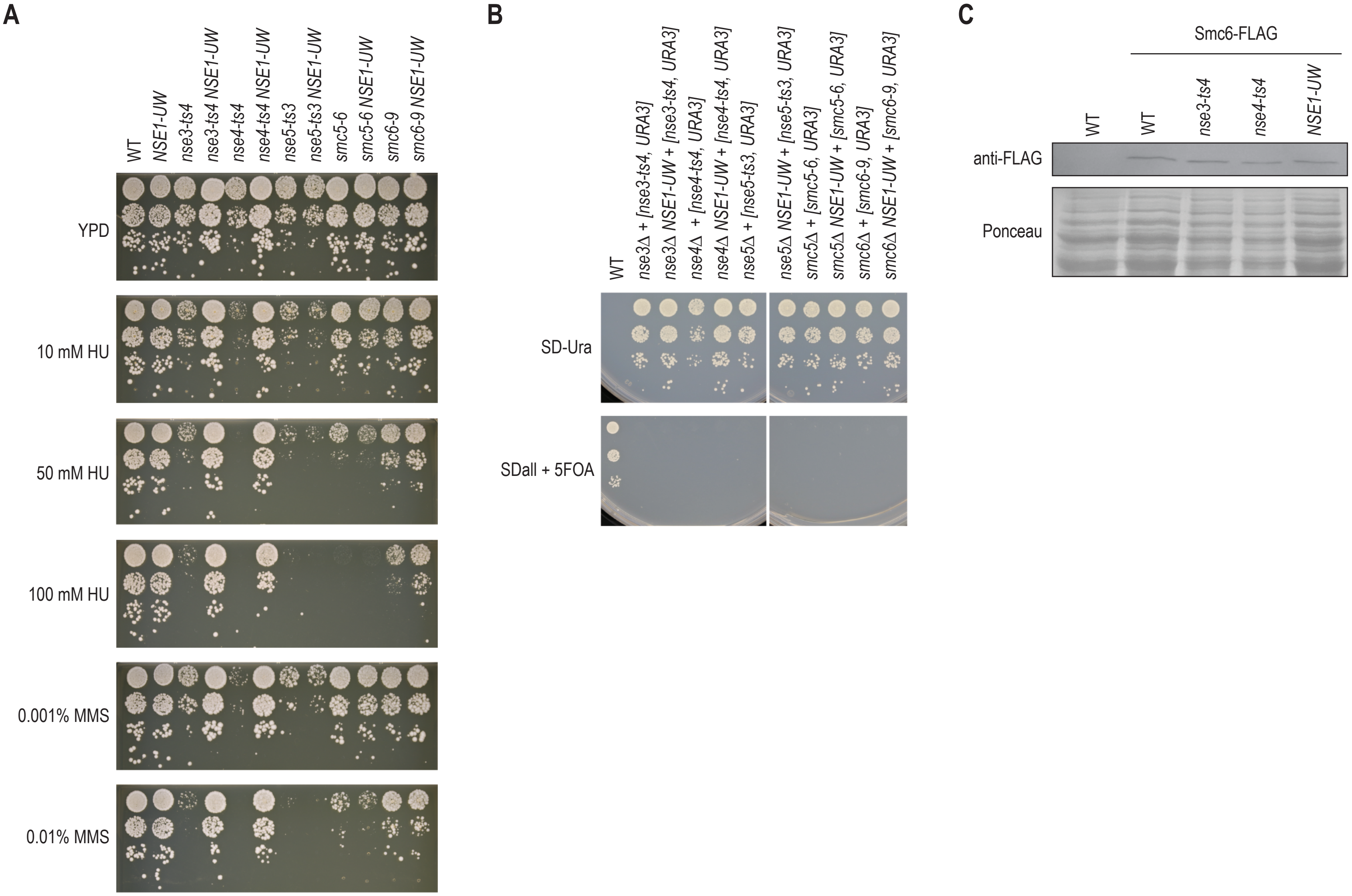


**Fig. S6. The *NSE1* allele of UWOPS87-2421 can suppress *NSE3* and *NSE4* TS mutants, but not deletion mutants.** (**A**) Suppression of the DNA damage sensitivity of *nse3-ts4* and *nse4-ts4* TS mutants by the *NSE1* allele of UWOPS87-2421. Cultures of the indicated strains were diluted to an optical density at 600 nm of 0.1 and a series of ten-fold dilutions was spotted on agar plates and incubated for 2-3 days at 30 ºC. UW = UWOPS87-2421, WT = wild type. (**B**) The *NSE1-UW* allele cannot suppress the lethality associated with deletion alleles of genes encoding SMC5/6 complex members. Spot dilutions were performed as described in (A). (**C**) Smc6-FLAG levels are not affected by mutations in *NSE1*, *NSE3*, or *NSE4*. Western blot analysis of Smc6-FLAG in the indicated strains. Ponceau staining was used as a loading control.
